# Biomaterial Degradation Products affect Regenerating Glia Independently of Surface Properties

**DOI:** 10.1101/2024.02.20.581223

**Authors:** Kendell M Pawelec, Jeremy ML Hix, Erik M Shapiro

**Affiliations:** Michigan State University, Dept Radiology, East Lansing, MI 48824; Michigan State University, Institute for Quantitative Health Science and Engineering (IQ), East Lansing, MI 48824

## Abstract

Devices to treat peripheral nerve injury (PNI) must balance many considerations to effectively guide regenerating nerves across a gap and achieve functional recovery. To enhance efficacy, design features like luminal fillers have been explored extensively. Material choice for PNI devices is also critical, as the determining factor of device mechanics, and degradation rate and has increasingly been found to directly impact biological response. This study investigated the ways in which synthetic polymer materials impact the differentiation state and myelination potential of Schwann cells, peripheral nerve glia. Microporous substrates of polycaprolactone (PCL), poly(lactide-co-glycolide) (PLGA) 85:15, or PLGA 50:50 were chosen, as materials already used in nerve repair devices, representing a wide range of mechanics and degradation profiles. Schwann cells co-cultured with dorsal root ganglion (DRG) neurons on the substrates expressed more mature myelination proteins (MPZ) on PLGA substrates compared to PCL. Changes to myelination and differentiation state of glia were reflected in adhesion proteins expressed by glia, including β-dystroglycan and integrin α_6_, both laminin binding proteins. Importantly, degradation products of the polymers affected glial expression independently of direct attachment. Fast degrading PLGA 50:50 substrates released measurable amounts of degradation products (lactic acid) within the culture period, which may push Schwann cells towards glycolytic metabolism, decreasing expression of early transcription factors like sox10. This study shows the importance of understanding not only material effects on attachment, but also on cellular metabolism which drives myelination responses.

## 1 Introduction

Peripheral nerve injury (PNI) remains a major healthcare burden worldwide [1,2]. Direct neurorrhaphy is the preferred treatment method, but excessive tension on the nerve precludes this approach as the gap increases beyond 10 mm [3]. Instead, surgeons implant grafts and devices to guide regenerating glia and neurons across the span. Unfortunately, as the distance between healthy proximal and distal nerve ends increases, treatment options become more and more limited.

The gold standard for functional repair remains the autograft, but this is in limited supply and causes donor site morbidity [4]. Decellularized allografts also possess the native architectural cues to guide regenerating axons, but they can have mismatches between host and donor nerves [5]. The limited supply of acceptable grafts has propelled the use of PNI devices. Clinically approved devices consist of hollow tubes that are secured to both ends of the injured nerve, and direct nerve regeneration across the gap. However, these simple devices result in limited functional recovery, and thus, alternative designs and features are in constant development [4].

There is general agreement that the key to a successful PNI device is providing guidance cues to the nerves while retaining 1) a high open lumen volume available for nerve ingrowth and 2) the ability to resist occlusion of the inner lumen due to kinking or movement around the implant. To satisfy these criteria, devices have been proposed with a variety of luminal fillers, porosity, and surface features [3,6]. In addition to structural features, a wide range of materials, or combinations of materials, have been investigated, consisting of both natural and synthetic polymers [4]. It is known that materials have a significant effect on the attachment and extracellular matrix deposition of glial cells [7,8]. Given the large differences in mechanical and chemical properties between the polymers used in PNI devices, we hypothesized that the substrate material also affects the nerve repair process.

To investigate, glia and neurons were co-cultured on microporous substrates, mimicking the regeneration process after injury. In this study, substrates were limited to synthetic polymers, as they offer advantages of mechanical strength that can be easily tuned with a wide range in degradation profiles. It was confirmed that the substrate polymer significantly affected nerve regeneration over multiple weeks, as measured by glial myelination markers. To dig deeper, Schwann cell attachment to the matrices and the effect of polymer degradation products were both examined. It was found that materials properties have a significant effect on the expression of glial attachment proteins, but that degradation products could affect glial marker expression independently of substrate attachment. Thus, material choice will play a role in the success of the implant technology independent of the overall device design.

## 2 Materials & Methods

Porous films were formed from polycaprolactone (PCL), poly(lactide-co-glycolide) (PLGA) 50:50 and PLGA 85:15. PCL (Sigma Aldrich); all had a molecular weight average of 80-90 kDa. PLGA 50:50 (Lactel/Evonik B6010-4) and PLGA 85:15 (Expansorb® DLG 85-7E, Merck) were both ester terminated. All three are biocompatible polymers used in clinically approved implants and have published usage as nerve implants [4,6,9].

### 2.1 Film manufacturing

Porous films were fabricated as substrates for in vitro studies, as previously described [10]. Polymers were solubilized in dichloromethane (DCM, Sigma); PCL was used at 4wt%, while PLGA was solubilized at 10wt% for comparable viscosity. Sucrose particles (mean particle size 31 ± 30 μm) were added to the solution as a porogen to create microporosity. The slurry, 70 vol% sucrose and 30 vol% polymer, was homogenized by vortexing for 10 minutes. Suspensions were cast and dried on glass sheets and subsequently submerged in deionized water. After 30 minutes in water to remove the sucrose, the microporous polymer film was air dried. In addition, for cell culture, non-porous PCL films were cast in the same way, without sucrose addition.

Similar surface morphology and properties of the films were verified via electron microscopy and contact angle measurements. To measure contact angle, films (porous and non-porous films that were cast in an equivalent manner but without addition of sucrose) were taped flat against a glass substrate. Distilled water was dropped onto the surface and the angle made with the surface was measured; reported measures are from 3 readings of 2 independent films each. Scanning electron microscopy was performed on a Zeiss Auriga, at 7keV in scanning electron mode, prior to sputter coating the samples with platinum.

### 2.2 Lactic Acid Concentration

To quantify the lactic acid concentration due to the degradation of polymer films during the culture period, disks of porous films (13 mm diameter) were punched from sheets. Prepared films were sterilized by immersion in 70% ethanol for 30 minutes, followed by two washes with sterile water. Films were placed into sterile microcentrifuge tube with 600 μl sterile distilled water, a volume equivalent the amount of complete media used during cell culture. Distilled water was used to remove noise and high background present in complete cell culture media. Degradation was conducted at 37°C and 5% CO_2_. Every 2-3 days until day 7, supernatant was removed and snap frozen on liquid nitrogen. To continue the degradation process, films were submerged in fresh water until day 7, a time point chosen to correspond to glial attachment studies. Distilled water incubated under the same conditions, but with no film present, was used as a blank media control.

Lactic acid analysis was performed on a Thermo Q-Exactive mass spectrometer coupled with a Thermo Vanquish UHPLC. Samples (in water) were prepared for LC-MS/MS analysis by adding 100 μl of sample to 900 μl acetonitrile. 5 μl of diluted sample was injected onto a Waters BEH-Amide UPLC column (2.1×100 mm) and separated according to the following 15 min binary gradient: initial conditions were 0% mobile phase A (90:10 water/acetonitrile containing 20 mM ammonium acetate and 0.1% ammonium hydroxide (from a 28-30% ammonia solution)) and 100% mobile phase B (90:10 acetonitrile/water containing 20mM ammonium acetate and 0.1% ammonium hydroxide (from a 28-30% ammonia solution)) and were held until 0.5 min, followed by a ramp to 50% B at 8 min, then a ramp to 20% B at 10 min, hold at 20% B until 11 min, then ramp back to 100% B at 11.1 min and hold until 15 min. The flow rate was 0.3 ml/min and column temperature was held at 40°C. MS data were acquired using electrospray ionization in negative ion mode with a capillary voltage of 2.5 kV, source defaults for 0.3 ml/min and S-lens RF level at 50. A data-dependent MS/MS method was used with survey scans acquired across m/z 70-1050 at a resolution setting of 35,000 3e6 AGC target and 100 ms max inject time. MS/MS scans (top 5) were acquired at 17,5000 resolution with an AGC target of 1e5, max inject time of 50 ms and an isolation window of 1.0 m/z. Data were processed using Excalibur software (Thermo Scientific). Both lactic acid and di-lactic acid (m/z 161.045) peak area was calculated. All data presented was run with 3 technical replicates.

### 2.3 Glial cell attachment

#### 2.3.1 Preparation of films

Prior to use, disks of porous films were punched (13 mm diameter) and sterilized as before. Sterile, hydrated films were placed into 24-well inserts (CellCrown™, Z681903-12 EA), as described previously [10]. Briefly, each sample consisted of two layers: a bottom layer of non-porous PCL (punched to 24mm diameter) and a top layer of porous film. After assembly into the inserts, they were placed at the bottom of a 24 well tissue culture plate, and the entire assembly was placed in UV light for 30 min.

Primary mouse Schwann cells, isolated from postnatal day 8 CD1 mouse nerves, were purchased (ScienCell Research Labs, M1700), and used before passage 5 in all experiments. Glia were passaged at 37°C and 5% CO_2_ in complete media (Dulbecco’s modified Eagle’s medium (DMEM, ThermoFisher Scientific, 11885), 10% fetal bovine serum (FBS, ThermoFisher Scientific, 160,000,036), 1% Penicillin– Streptomycin (ThermoFisher Scientific, 15,140,122), 21 μg/ml Bovine Pituitary Extract (BPE; Lonza, CC-4009), and 4 μM forskolin (Calbiochem, 344270)). Unless noted, sterilized substrates were coated with laminin (natural mouse laminin, Invitrogen (23017-015)). For coating, each sample was incubated with 4 μg of laminin, diluted in sterile water, for 1 hr at room temperature. The surfaces were then washed and left in sterile water, at room temperature until used for cell culture, within 2 hr of coating. For attachment studies, cell number was quantified at 24 hrs post-seeding. To analyze attachment proteins, samples were harvested at day 7. Samples were fixed for immunohistochemical staining (IHC) at day 1 and 7.

#### 2.3.2 Attachment

Cell number at 24 hrs post-seeding was quantified on both untreated or laminin coated substrates. On each film, 1×10^4^ cells/well were plated in 400 μl of either complete media or control media (without FBS). Samples were harvested 24 hr after seeding, by washing in phosphate buffer saline (PBS) and storing films at −80 °C for DNA quantification using Quant-iT™ PicoGreen™ dsDNA Assay Kit (ThermoFisher Scientific), per manufacturer’s instructions. Frozen inserts were digested in 200 μl papain buffer (0.1 M phosphate buffer, 10 mM L-cysteine, 2 mM Ethylenediaminetetraacetic acid (EDTA), 3 U/ml papain) at 60 °C overnight. In the assay, samples were diluted 1:4 in TNE buffer and loaded on a black 96 well plate. An equal amount of PicoGreen dye (1:200 in TNE buffer) was added to each well, and fluorescence was read at Ex: 480 nm, Em: 520 nm using a BioTek plate reader. Results were compared to a standard of Schwann cells (0–5×10^4^ cells), created on the day of experimental cell seeding. All tests were done in triplicate, with two technical replicates each, and presented as mean ± standard error.

#### 2.3.3 Direct and indirect culture

For analysis of attachment proteins when cells were adhered to porous films, 5×10^4^ cells/well were plated onto inserts in complete media, and cultured for one week, changing media three times per week. To investigate the indirect effects of the porous films, 2-3 days prior to culturing cells, porous disks were incubated in 600 μl complete media at 37°C and 5% CO_2_; as a control, 600 μl media was incubated without a film. Cells were seeded onto laminin coated wells of a 24 well plate and allowed to attach in complete media for 8 hrs. After, the supernatant was removed and replaced with 500 μl supernatant incubated with porous films (or no film control). Fresh media was added to the soaking films to be used at the next media change (3 days later). At day 7, all remaining samples were analyzed for protein expression; 3-4 biological replicates, with 2 technical replicates each.

### 2.4 Co-culture model

The co-culture model simulating nerve regeneration was adapted from previously published literature [7].

#### 2.4.1 Animal care

All animal handling and surgical procedures were performed in compliance with local and national animal care guidelines and approved by Michigan State University’s IACUC. For all experiments, 4 week-old CD1 female mice were used (Charles River Laboratories). Mice were group-housed in a 12-h light-dark cycle and provided with enrichment environment (EE) (e.g. nestlets, plastic huts); rodent chow (Teklad Global Diet® 2918) and water (facility tap) was offered ad libitum.

#### 2.4.2 Preparation of dorsal root ganglion (DRGs) cultures

Naive DRGs at all rostro-caudal levels along the spinal cord were harvested immediately following euthanasia via CO_2_ inhalation. For each biological replicate, DRGs from 3-4 individuals were combined. DRGs were harvested, placed in ice-cold Leibovitz’s L-15 medium (Gibco), and digested in Collagenase type II (4 mg/ml, Worthington) and Dispase II (2 mg/ml, Sigma), diluted in L15 at 37 °C, for 45 min. Tubes were inverted every 10 min during enzymatic digestion. The digestion medium was pulled off and cells were rinsed with neural growth medium (DMEM/F-12/ GlutaMAX + 10% FBS + B27 supplement + Penicillin/Streptomycin).

#### 2.4.3 Co-culture model

Porous films were coated with laminin prior to use, as before. Co-cultures consisted of seeding 1×10^4^ Schwann cells on each film, followed by the addition of 6-8×10^3^DRG neurons per well, 4–6 h after initial Schwann cell seeding. During the first 24 h, the inserts were cultured in complete DRG medium. Following this, co-cultures were maintained for 3 days in Schwann growth medium (DRG complete medium, 21 μg/mL BPE, 4 μM forskolin). Finally, the culture medium was replaced with differentiation medium (DRG complete medium, 50 mg/ml ascorbic acid, 21 μg/ mL BPE, 0.5 μM forskolin). Media was aspirated and replaced three times per week for up to 2 weeks. Samples were taken at 1 week for immunohistochemical staining (IHC) and weeks 1 and 2 for analysis of protein expression.

### 2.5 Protein Expression

To extract proteins, inserts with cells were washed once in PBS and placed in Ripa buffer (Fisher Scientific) with 1× protease and phosphatase inhibitor cocktail (Sigma, PPC1010): 100 μl for glia alone and 200 μl for co-cultures. For harvest from tissue culture plates, 100 μl of lysis buffer was added directly to washed wells. Cells were scraped from the substrate and the buffer was then transferred to a microcentrifuge tube. Samples were rocked gently overnight at 4 °C; the inserts were then removed and the protein lysate was stored at −20 °C until use.

Protein concentration was measured (BioRad, BCA kit) prior to quantifying expression of myelination and attachment markers using sodium dodecyl sulfate polyacrylamide gel electrophoresis (SDS-PAGE) and Western blotting. Full protocol details and images of complete blots are presented in Supplemental. Briefly, samples with 2× Laemmli buffer were incubated at 60°C for 20 min and separated by SDS/ PAGE. After transfer to PVDF membranes (iBlot 2 Gel Transfer Device, invitrogen), membranes were blocked with 5% dried milk in PBST (phosphate buffered saline pH 7.4, containing 0.1% Tween-20) and probed with primary antibodies specific for Oct-6 (1:1,000, Aviva Systems Bio, ARP33061_T100), MPZ (1:1000; Aves, PZO), c-Jun (1:1000, Cell Signaling Technology, 60A8), Sox10 (1:1000, Abcam, ab155279), integrin α_6_ (abcam, ab181551), integrin β1 (abcam, ab179471), paxillin (1:1000, BD Biosciences, 612405), β-dystroglycan (1:1000, Santa Cruz, sc-165997), and integrin α_V_ (1:1000, abcam, ab302640) at 4 °C overnight. The next day, the membrane was washed and incubated with anti-chicken IgY-HRP (Millipore, 12-341), anti-mouse IgG-HRP (Millipore, 12-349), or anti-rabbit IgG-HRP (abcam, ab205718) at room temperature for 2 hr. Protein bands were visualized with Amersham ECL Plus substrate (cytiva, RPN2232) on a Li-Cor Odyssey FC and resulting signal was quantified via Image J.

Developed membranes were washed in PBS and stripped for 15 minutes in Restore Western Blot Stripping Buffer (Thermo Scientific 21059) before being either blocked and reprobed, or stained for total protein using the BLOT-Fast Stain (G Biosciences, cat# 786-34) according to manufacturer’s directions. The total protein signal was imaged under visible light on a c300 imager (azure biosystems). Protein expression was quantified via Image J and normalized to total protein to adjust the measured band intensity. For comparisons between protein expression from cells cultured directly on substrates, all signals were normalized to PCL, which was run on every membrane. For comparison of proteins in indirect culture, signal was normalized to blank (control) samples. All data reported are the result of 3-4 independent replicates (two technical replicates each) and are reported as mean ± SEM. Images of all blots are presented in Supporting Information.

### 2.6 Immunohistochemistry

Samples were fixed in 4% paraformaldehyde for 15 min and stored in PBS at 4 °C, prior to immunohistochemical staining (IHC). Samples were washed in PBS+0.1% Tween-20 (PBST), then permeabilized in 0.1% Triton X-100 in PBS, followed by two PBST washes. Samples were then blocked with 5 wt% bovine serum albumin (BSA) in PBST, for 1 hr, at room temperature. After washing twice in PBST, films were incubated with primary antibodies, overnight at 4 °C, in 3 wt% BSA/PBST. Following primary incubation and two PBST washes, secondary antibodies in PBS were incubated with the samples for 1–2 h at room temperature. If the actin cytoskeleton was visualized, samples were incubated for 30 minutes in ActinRed™ 555 ReadyProbes™ (Invitrogen, R37112, 2 drops per ml PBS). Finally, samples were washed twice with PBS and left in a solution of PBS and DAPI stain (1:000 in PBS, ThermoFisher Scientific, 62248). Primary antibodies, and the dilutions used, were: β-tubulin (1:1500, TUJ1, Promega, G7121), sox10 (1:300, Abcam, ab155279), GFAP (1:500, DAKO, Z033429; EMD Millipore, AB5541), paxillin (1:500, BD Biosciences, 612405). Secondary antibodies: Alexa Fluor 488 (1:1000, A11070), Alexa Fluor 647 (1:1000. A21236), Cy3-AffiniPure Donkey Anti-Chicken IgY (1:500, Jackson ImmunoResearch, 703-165-155).

Stained films were inverted onto a coverslip in a drop of Slow Fade Diamond Antifade Mountant (Invitrogen, S36972). Imaging was performed using a Leica DMi8 microscope, with an LASX software interface. With the high surface roughness of the films, z-stacks were taken at a minimum of 3 places on the film and post-processed with Thunder image analysis (Leica) to remove background fluorescence.

### 2.7 Statistics

All statistics were done using GraphPad Prism 10.1.2. Data was first analyzed via ANOVA, followed by multiple comparisons of the means, using Fisher’s LSD test. In all cases, α < 0.05 was considered significant, with a 95% confidence interval.

## 3 Results

Polymer substrates regulate glial response, which could have short- and long-term consequences for PNI devices. The three polymer matrices tested all had similar wettability (80-90 degrees), which was dominated by the addition of microporosity within the substrate, Figure 1(a), dropping significantly. While films had equivalent surface characteristics and could support the attachment and growth of glia and neurons, Figure 1(b-d), the polymer matrix significantly affected glial markers and myelin protein expression at one and two weeks of culture, Figure 1(e-f). Glia cultured on PCL had significantly higher expression of sox10, cJun and GFAP, all markers for an immature or repair phenotype after nerve damage. On PLGA substrates, PLGA 50:50 in particular, glia tended to express greater levels of later stage markers of myelination, like MPZ.

**Figure 1.**
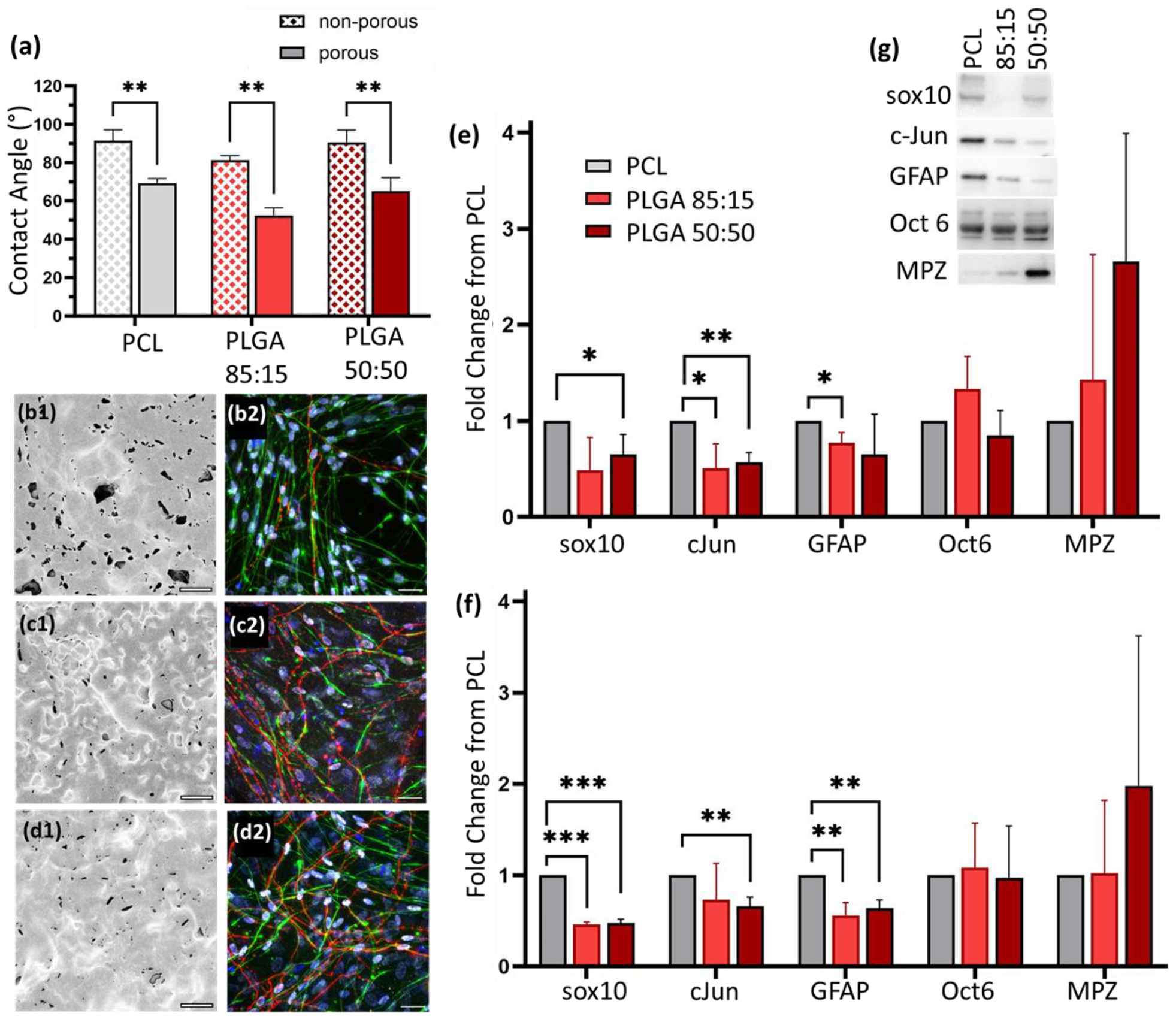
Matrix polymer significantly affects expression of myelination markers in co-cultures of glia and DRG neurons. Polymer films had similar (a) wettability for porous and non-porous films and (b1-d1) surface morphologies, visualized via electron microscopy: (b) PCL, (c) PLGA 85:15 and (d) PLGA 50:50. (b2-d2) Glia and neurons could adhere and proliferate on polymer substrates after one week in culture on all substrates; red: neurons (TUJ1), green: Schwann cells (GFAP), white: sox10, blue: nuclei (DAPI). Protein expression of glia at (e) 1 week and (f) 2 weeks showed persistent changes across three different matrices; all data is reported as a fold change from PCL and protein loading is normalized by total protein staining. (g) Representative protein bands from Western blotting at week 1. Data presented as mean ± SEM. Scale bar (b-d): 25 μm. * p< 0.05, ** p< 0.01, ***p<0.001.

### 3.1 Glial Attachment

The polymer matrices all supported glial attachment with no significant difference in percentage attachment when cells were seeded onto substrates with or without coating with laminin (LN), Figure 2(a). Only addition of FBS to the culture media significantly increased glial attachment. Comparing the attachment of different polymer substrates, trends were visible even without coating the substrates with laminin (LN) or the addition of FBS to media, Figure 2(a). Attachment reached 19.61 ± 11.9% and 23.27 ± 11.9% for PCL and PLGA 85:15, respectively. In all cases, the poorest attachment occurred on PLGA 50:50, reaching a maximum of 3.95 ± 0.5%. While the difference became significant only after coating the surface, the consistent trend makes this unlikely to be a property of protein binding/corona on the surfaces of the polymer. In all cases, attachment of glia at 24 hours shows elongated and spread cells, Figure 2(b-d).

**Figure 2.**
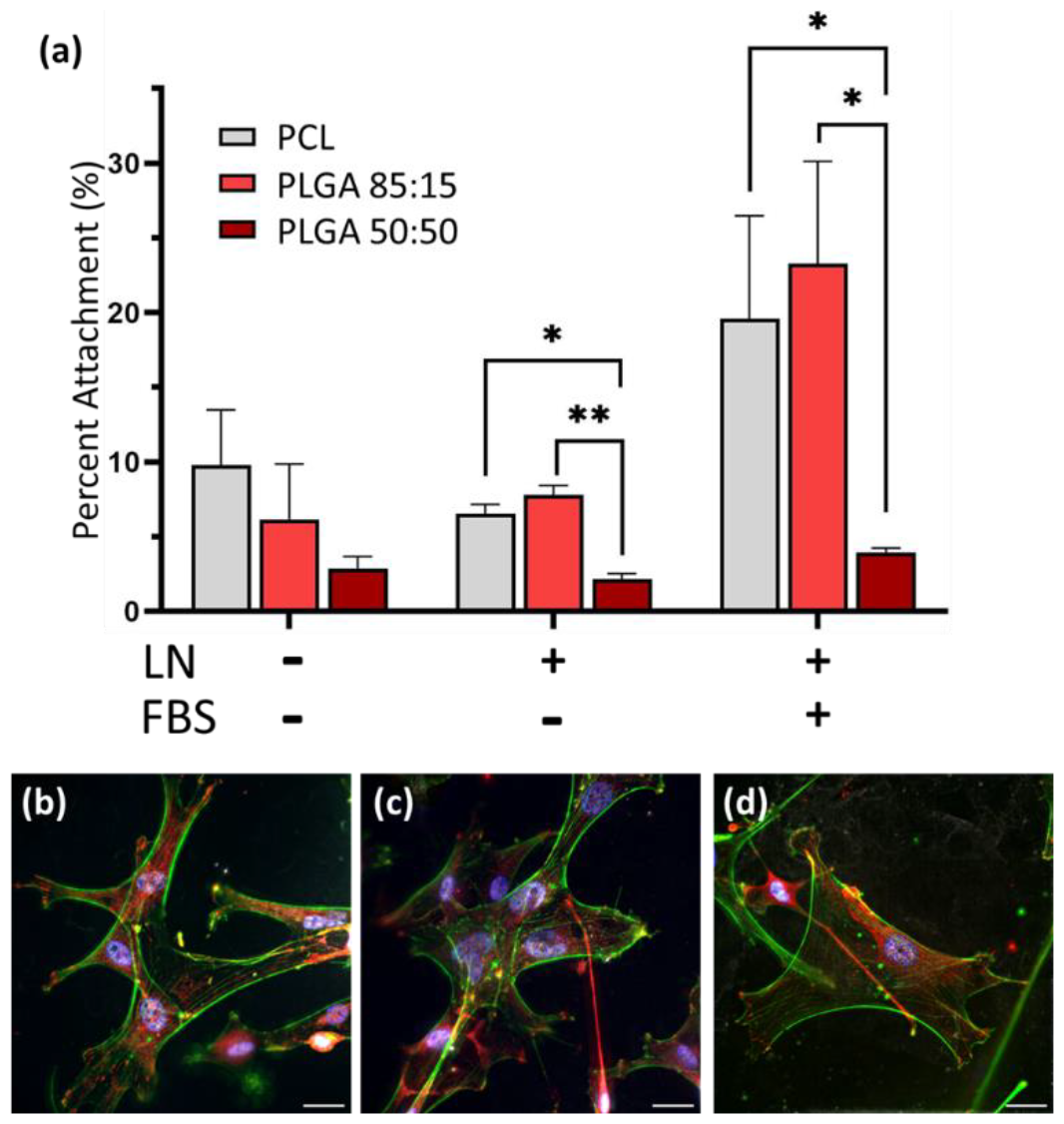
Trends in the attachment of glia were not significantly altered with protein adsorption. (a) At 24 hours post-seeding, the percent attachment ranged from 5-10% without laminin (LN) coating or FBS present in the media. Laminin had no significant effect on average attachment, while FBS significantly increased attachment. Glial cells, cultured on LN coated films, were elongated on all microporous films and expressing glial markers (sox10): (b) PCL, (c) PLGA 85:15 and (d) PLGA 50:50. Blue: nucleus (DAPI), white: sox10, green: actin, red: paxillin. Scale bar (b-d): 25 μm. Data presented as mean ± SEM. * p< 0.05, ** p< 0.01.

As cellular attachment was affected by polymer substrate, expression of attachment proteins also varied on different matrices, Figure 3. Laminin binding proteins, including integrin α_6_ and β-dystroglycan, were most markedly affected by matrix polymer and were present from initial glial attachment at day 1 through a week in culture, Figure 3(a-c). Glia cultured without neurons had an increased protein expression of β-dystroglycan on PLGA substrates. When cultured in contact with neurons, the most significant changes in attachment occurred with a down-regulation of integrin α_6_ on PLGA 50:50. Attachment integrins present in Schwann cell focal adhesions, such as paxillin, and those which bind primarily via RGD motifs not present on laminin, including integrin α_V_, were not affected overall [11].

**Figure 3.**
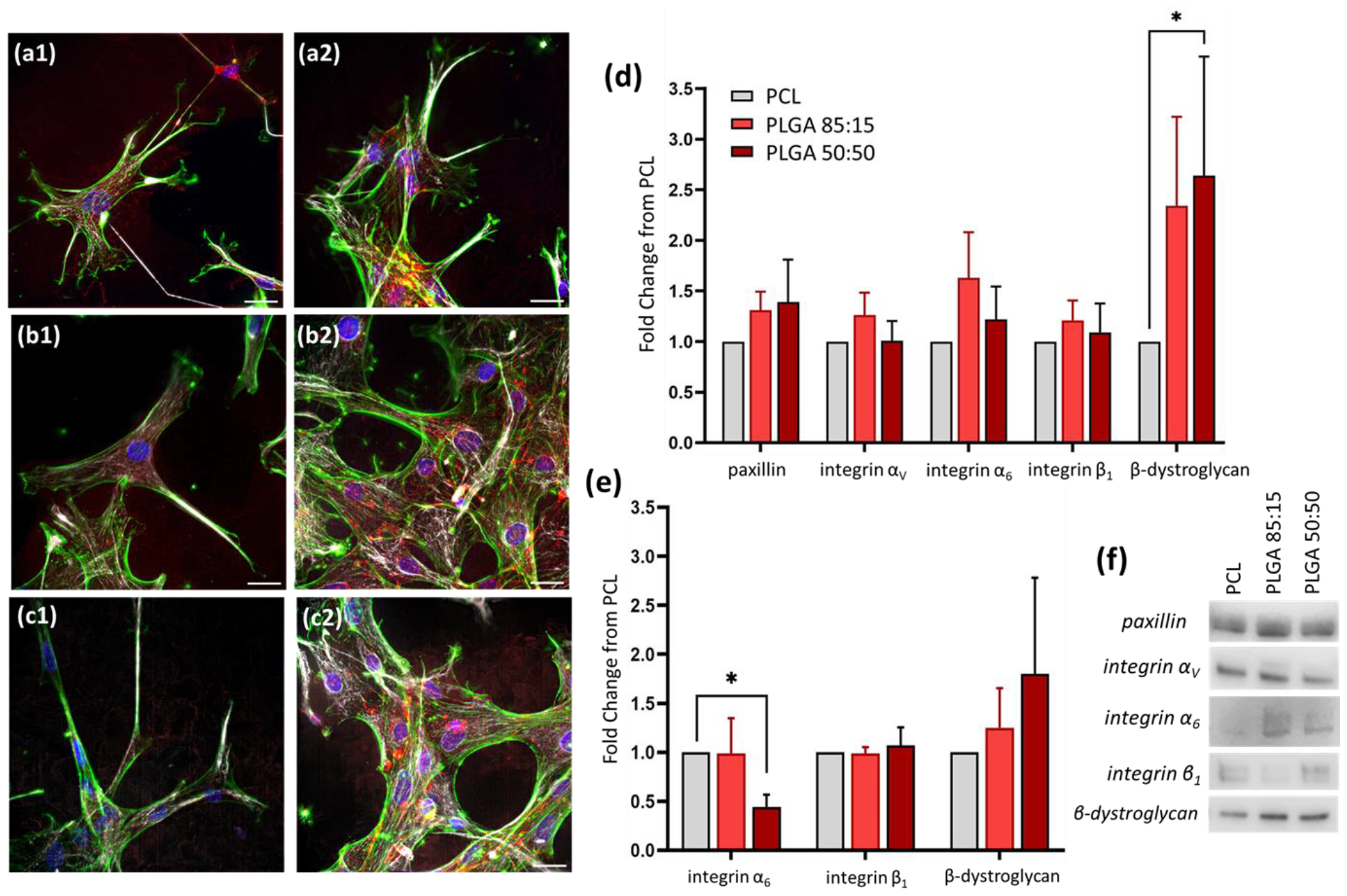
Attachment proteins that recognize laminin were significantly altered by polymer substrate. Glia were cultured for (1) 24 hrs and (2) 7 days on different matrices (a) PCL, (b) PLGA 85:15, (c) PLGA 50:50. All glia express laminin attachment proteins integrin α_6_ (white), and β-dystroglycan (red); blue: nuclei (DAPI), green: actin. Expression of glial attachment proteins at week 1 for (d) glia alone and (e) glia in co-culture with neurons. (f) Representative protein bands for attachment proteins. All protein expression is reported as fold change from PCL and loading was normalized by total protein on each blot. Data presented as mean ± SEM. Scale bar (a-c): 25 μm. * p< 0.05.

### 3.2 Effect of Degradation Products

Degradation products from the polymer matrices affect glial markers, independently of substrate attachment. Indirect culture of glia with media incubated with polymer substrates showed significant differences in glial markers, notably sox10, Figure 4. However, the change in glial markers was not linked to differences in substrate attachment. Differences in glial marker expression mirror those observed in co-cultures.

**Figure 4.**
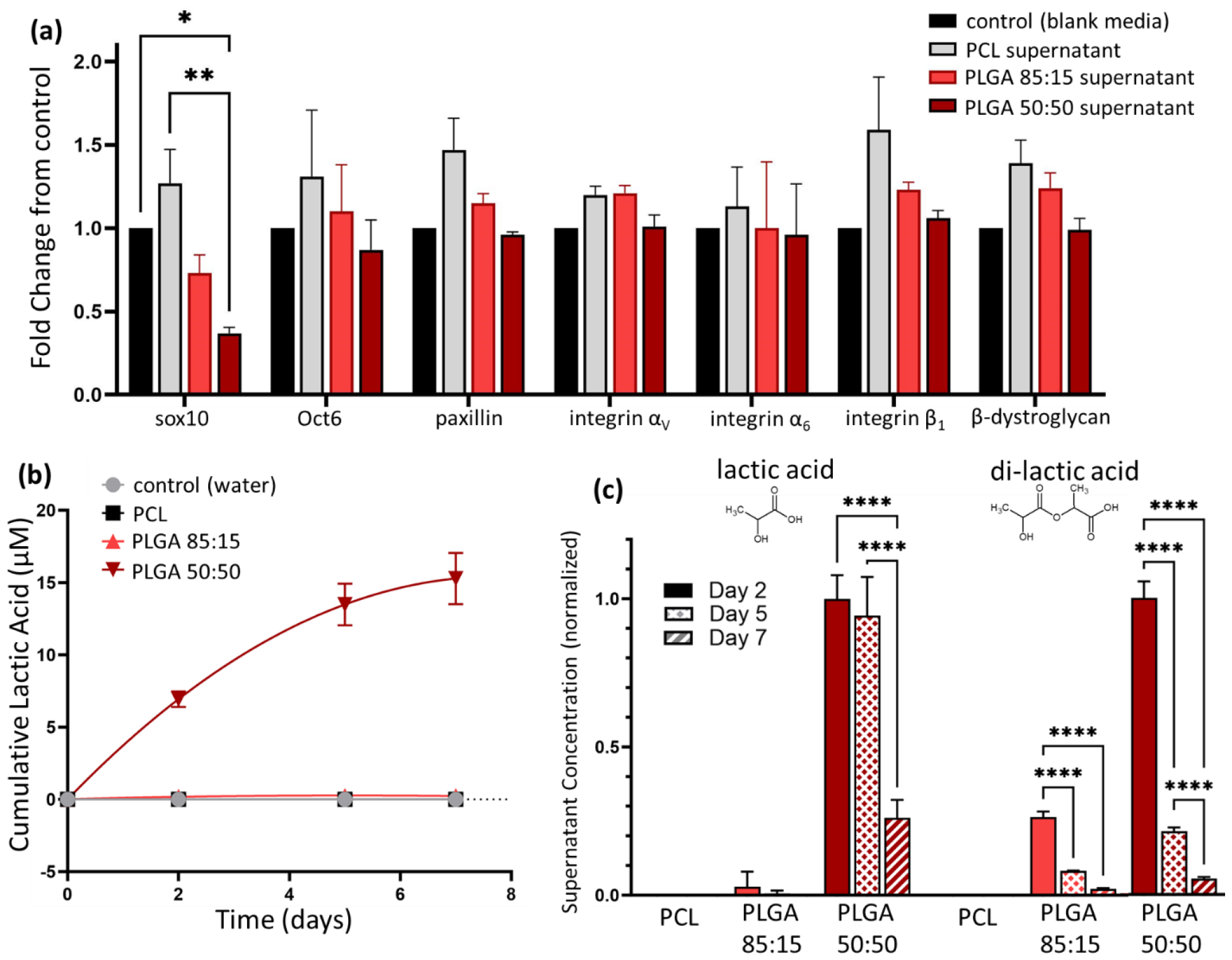
Indirect culture of glial with polymer matrices affected glial markers after one week. (a) Expression of attachment proteins was not altered, but glial marker sox10 was deferentially expressed, following trends seen in co-cultures. All data is reported as fold change from control samples incubated in media that was not in contact with a polymer matrix. Protein loading was normalized by total protein staining. Data presented as mean ± SEM. Lactic acid concentration was quantified over one week of polymer degradation. (b) Cumulative release of lactic acid was only visible in PLGA 50:50. (c) Both PLGA 50:50 and PLGA 85:15 had measurable amounts of di-lactic acid present in extract media; data was normalized by PLGA 50:50 on day 2. Lactic acid data (b-c) is presented as mean ± standard deviation. * p< 0.05, ** p< 0.01, **** p<0.0001.

Polymer degradation extracts were further analyzed over one week for levels of lactic acid, a key degradation product of PLGA films and a known biological signaling molecule [12]. Extracts of PLGA 50:50, the fastest degrading polymer, contained measurable lactic acid by day two (6.94 ± 0.6 μM), with a cumulative release of 15.28 ± 1.8 μM over one week. No other samples had significant release of lactic acid. However, both PLGA 50:50 and 85:15 had measurable amounts of di-lactic acid, Figure 4(c).

## 4 Discussion

Devices to repair PNI are specialized to guide regenerating neurons and their supporting glia to bridge damaged tissue. However, the hollow tubes available in the clinic fall short of robust functional regeneration [4]. In redesigning peripheral nerve devices, engineers have incorporated features like microporosity and tube fillers, such as linearly aligned channels. Material choice in medical devices must also be considered carefully, as material properties can affect biological response through direct contact, via adhesion mechanisms [13], and indirectly through their degradation products [14]. While natural polymers contain biocompatible motifs and can be naturally broken down by enzymes and cells, they lack mechanical strength. For this reason, and for their tightly controlled chemistry, synthetic polymers are becoming the material of choice for medical devices.

Of the biocompatible synthetic polymers available, PCL and PLGA have a long history of use as biomedical devices. Both have demonstrated compatibility for nerve regeneration, having been shown to successfully support axon regrowth in multichannel devices [6,15]. While both PCL and PLGA are polyesters, they have very different chemistries, and therefore materials properties, such as surface charge, mechanics and susceptibility to hydrolyzation. These materials properties are all important for determining the biological response and can affect PNI device design [9], which necessitates more in-depth study.

To better understand the range of biological response to synthetic polymer matrices, the macroscopic properties of the substrate films in this study were kept as similar as possible. Thus, all films are 70vol% microporous, with equivalent surface roughness [10]. From macroscopic measurements, polymer films were all hydrophobic (90-degree contact angle). The addition of microporosity, altered wettability, creating more hydrophilic surfaces, with contact angels between 52-69 degrees. While wettability and surface topography were similar, mechanics and degradation were not. PCL is more compliant than PLGA and more resistant to hydrolysis, requiring years to degrade in physiological environments [10,16]. Further, two PLGA blends, differing in the ratio of poly(lactic acid) to poly(glycolic acid) were examined in this study. While the blends have very similar mechanics [10], a higher content of hydrophilic glycolic acid is known to increase degradation rate [17].The two blends allowed investigation into how the rate of material hydrolysis (ie degradation) affected biological response.

Following literature, all polymer matrices supported the regeneration of nerve tissue, measured by markers of myelination, in a co-culture model of glia and DRG neurons. Significant differences in glial markers were noted between PCL and PLGA substrates, and these differences remained significant over 2 weeks, suggesting a long-term effect of the substrate used for PNI devices. PCL substrates appeared to support a higher proportion of repair glial cells, those that have dedifferentiated after nerve damage and are still in early stages of turning on pro-myelination machinery. This is consistent with the upregulation of early transcription factors like c-Jun and GFAP related to initial dedifferentiation and repair [18]. The changes in biological markers of myelination follow the differences in Schwann cell adhesion observed across different substrate polymers [7,8,10]. This raised the question of whether differences due to polymer matrices were caused by direct mechanisms, such as those mediated via attachment and adhesion proteins, or via indirect exposure to degradation properties.

### 4.1 Glial Attachment

Cellular attachment is a complex process that evolves over time. Many surface properties of materials are known to affect the efficacy of cellular attachment, including surface topography, hydrophobicity, chemistry, and protein adhesion. Addition of microporosity, to mimic surfaces in use for nerve repair [6], increased surface roughness and dominated attachment characteristics of the substrates. It has previously been shown that surface coatings with adhesion molecules do not affect attachment on rough films, while significant changes are observed on flat substrates of the same material [7]. Surface roughness also has greater affect than coating protein on cellular morphology [7].

Glial cell attachment is affected by polymer substrates [10]. These differences were observed in the present study as well. In particular, PLGA 50:50 had low Schwann cell attachment. This low attachment was not related to adsorption of an adhesion protein (laminin) at the surface or affected significantly by FBS in the culture media. In contrast, PLGA 85:15 had the highest glial attachment, particularly in the presence of FBS. Macroscopically, the wettability of the films was not altered by polymer blend, but at the nanoscale, the chemical side chains presented to cells may be different and the faster hydrolyzation of PLGA 50:50 could also affect cellular response.

As expression of adhesion proteins has been shown to be crucial to cellular attachment and activity in a variety of systems [19], changes in expression of attachment proteins were investigated. The basal lamina of Schwann cells contains several key matrix components, including laminin and collagen Type V [20]. Therefore, glia have the ability to recognize and bind to different motifs. Fibronectin and gelatin have RGD binding sites that are recognized by several integrins, including α_V_, and these play a role in Schwann cell spreading in vitro [11].

Here, substrate polymer did not affect expression of RGD binding integrins. Focal adhesions have also been reported to be involved in Schwann adhesion, and paxillin, a protein within the focal adhesion complex, is critical for Schwann cell motility [21]. Again, expression remained consistent across different substrates. Instead, the laminin coating on the polymer films dominated the attachment mechanisms investigated. Only attachment components known to be associated with laminin binding were differentially expressed, including β-dystroglycan [22] and integrin α_6_ [23]. This substrate coating was chosen to mimic conditions present in co-culture models, where laminin is required for axonal growth [7].

When glia were cultured without neurons present, cells expressed higher levels of β-dystroglycan on PLGA surfaces. When Schwann cells were grown in contact with neurons, the expression of laminin binding components changed. PLGA 50:50 had downregulated integrin α_6_, although expression of β-dystroglycan remained elevated. Laminin receptors have distinct roles during growth and myelination, with altered expression depending on glial state [23]. Integrin α_6_is the attachment protein available at all stages of Schwann growth and differentiation, while β-dystroglycan is expressed at the cell membrane just prior to myelination. This supports the hypothesis that PLGA substrates tend to support Schwann cells in a more mature pro-myelinating phenotype compared to PCL substrates, but this did not answer the question of whether changes were due to direct contact alone or affected by degradation products as well.

### 4.2 Degradation Products

Degradation products of polymers are oligomers and monomers that diffuse away from the device. In vivo, degradation products generally remain close to the site of implantation, slowly diffusing via the blood to the kidneys and liver [24]. Degradation products can have alternate activity in the body and be potentially toxic after release [25]. A major advantage to PLGA has historically been its highly biocompatible degradation products, lactic acid and glycolic acid, that can be removed by feeding into gluconeogenesis or into the tricarboxylic acid cycle for ATP production [12]. However, in vivo, may of the intermediates or substrates of metabolic pathways have additional roles. Lactate, in the form of a dissociated lactate anion from lactic acid, is one such metabolite that acts as a multifunctional signaling molecule within the body in various tissues [12]. It has particularly important roles in homeostasis of nerves in the central and peripheral nervous system [26].

The effect of degradation products was measured on cultures of glial cells without neurons or pro-myelinating media, to ensure the changes in expression were due to glia alone. Indirect culture with soluble degradation products did not affect the attachment of glial cells. Modification of cellular attachment is most often associated with changes to the substrate’s chemistry, mechanics or topography [27], which remained unchanged across sample groups. However, expression of the pan-glial marker, Sox10, was affected by degradation products, mimicking the expression pattern seen in a myelinating co-culture system in direct contact with polymer films. Thus, the relative expression of glial markers was, at least in part, mediated by a soluble factor from polymer degradation. PLGA 50:50, the film with the highest degradation rate, produced the strongest change in expression and released the greatest amount of lactic acid. This has been noted in the immune system, were even at short time points (<12 days), polymers are capable of releasing oligomeric degradation products that alter the cellular response via a switch to glycolytic metabolism [14,28]. This study confirmed that measurable lactic acid was released by PLGA 50:50 even by day 2, and that oligomeric products were present in all PLGA blends. For comparison, the cumulative release of lactic acid by PLGA 50:50 was around 15 μM over one week, compared to baseline measures of lactic acid in complete media which are between 20-30 μM.

Myelinating Schwann cells are known to be sensitive to lactate concentration, and express transporters for lactate that are linked to myelination potential [29]. With increased lactate present in culture medium, Schwann cells have greater expression of transcription factors that drive myelination and upregulate of myelin protein mRNA [29]. This supports the finding that PLGA 50:50, and to a lesser extent PLGA 85:15, drives glia to express greater amounts of mature myelin proteins (MPZ), and showed alteration in binding mechanism, as dystroglycan is linked to a myelinating cell phenotype compared to an immature glial phenotype. Surface characteristics must still play a role in attachment protein expression. It was only in direct contact with polymers that attachment machinery was significantly altered, and not via degradation product expression. Therefore, surface adhesion and degradation products affect cellular response independently, although the pathways may act synergistically to determine Schwann cell fate and myelination potential.

Putting these results in a clinical context requires balancing not only the biological response to polymer selection, but practical considerations of nerve repair devices including implantability and long-term performance. While PLGA 50:50 promotes the greatest myelination response in glia, the degradation rate may be too fast for the support of nerve regeneration across a clinically relevant gap of up to 3 cm. Also, PLGA is much more brittle when compared to PCL, and devices utilizing PLGA would be prone to permanent deformation and kinking if used in areas that experience high movement [9]. Certainly, more attention should be paid to the potential effects of biomaterials on the glycolytic metabolism in the PNS when creating nerve repair devices.

## 5 Conclusions

Material choice for PNI devices can have both acute and sustained impact on nerve repair. This was demonstrated with the support of a more mature myelination profile of glial cells on PLGA substrates as opposed to PCL over 2 weeks in culture. Adhesion of Schwann cells was affected by the polymer substrates, with significant changes in laminin binding proteins of both glia alone and glia in co-cultures. Again, higher expression of β-dystroglycan on PLGA pointed to a more mature pro-myelination phenotype in Schwann cells. Measurable degradation products, namely lactic acid, were present in the supernatant of PLGA 50:50 substrates even within the first week of culture. As lactate is a mediator for glycolytic metabolism in the body and very important for nerve tissue homeostasis, it had a measurable effect on Schwann glial markers, independent of substrate attachment. Therefore, in balancing properties for nerve repair devices, the choice of material should not be an afterthought.

## 6 Acknowledgments

The authors would like to thank the Michigan State University (MSU) Mass Spectrometry and Metabolomics Core for performing analysis of polymer extracts. This study was supported by the National Institute of Biomedical Imaging and Bioengineering of the NIH under award number R01EB029418. The content is solely the responsibility of the authors and does not necessarily represent the official views of the National Institutes of Health.

## Notes

### Competing Interest Statement

The authors have declared no competing interest.

