## Supplemental for "Biomaterial Degradation Products affect Regenerating Glia Independently of Surface Properties"

### S.1 Protein Expression

Western blotting was used to quantify protein expression. Co-cultures were lysed in Ripa buffer (supplemented with phosphatase inhibitors and protease inhibitors), and spun down at 15,000×g for 5 min at 4 °C. After which, supernatants were collected and the protein concentration of the supernatant was measured (BioRad, BCA kit). Supernatants were combined with 2× Laemmli buffer, incubated at 60°F for 20 min, separated by SDS/ PAGE (4 µg of protein per lane), and transferred to PVDF membranes. Transfer was accomplished using the iBlot 2 Gel Transfer Device (invitrogen, IB21001), a dry transfer system, at the recommended setting of 20 V for 1 min, 23 V for 4 min and 25 V for 2 minutes (7 min total). PVDF membranes were blocked with 5% dried milk in PBST (phosphate buffered saline pH 7.4, containing 0.1% Tween-20) and probed with primary antibodies at 4 °C overnight. For co-cultures, antibodies were specific for Oct-6 (1:1,000, Aviva Systems Bio, ARP33061_T100), MPZ (1:1000; Aves, PZO), c-Jun (1:1000, Cell Signaling Technology, 60A8), Sox10 (1:1000, Abcam, ab155279). For glial attachment and effect of degradation products, additional primaries were integrin $\alpha_6$ (abcam, ab181551), integrin $\beta_1$ (abcam, ab179471), β-dystroglycan (1:1000, Santa Cruz, sc-165997), paxillin (1:1000, BD Biosciences, 612405), integrin $\alpha_V$ (1:1000, abcam ab302640). The next day, the membrane was washed and incubated with anti-chicken IgY-HRP (12-341, Millipore) or anti-rabbit IgG-HRP (12–348, Millipore, ab205718, abcam) at room temperature for 2 hr and further washed in PBST six times. Early problems with dark spots on developed membranes were traced to agglomeration of the secondary anti-rabbit IgG-HRP (12–348, Millipore) after storage; from that point, the anti-rabbit IgG-HRP ab205718 (abcam) was used exclusively.

Protein bands were visualized with ECL Plus substrate (cytiva, RPN2232) on a Li-Cor Odyssey FC and resulting signal was quantified via Image J. Membranes were then washed in PBS and stripped for 15 minutes in Restore Western Blot Stripping Buffer (Thermo Scientific 21059) with subsequent PBS washing. Membranes were either blocked and reprobed or were stained for total protein using the BLOT-Fast Stain (G Biosciences, cat# 786-34) according to manufacturer's directions. The total protein signal was imaged under visible light on a c300 imager (azure biosystems).

Signal for total protein in each lane was quantified on Image J, and the protein loading was normalized to the signal from the first lane. The normalized signal was used to adjust the measured band intensity from antibody staining, also quantified using Image J. For comparison, all protein expression was recorded as a fold change from PCL matrices. The final protein quantification reported for co-cultures is the result of three biological replicates (2 technical replicates each). For glial culture, quantification is the results of five biological replicates (2 technical replicates each); due to limitations of protein content not all antibodies were assayed for all replicates.

In the following figures, each set of membranes is presented with all protein bands shown, along with the total protein staining for each membrane. Where a molecular weight marker is not present on the blot, the molecular weight of bands was determined from an initial blot where the molecular weight marker was present. For integrin $\alpha_6$ expression, only bands at high molecular weight, corresponding to the mature protein, were quantified. For β-dystroglycan only the 43kDa band was quantified, as that is the active form.

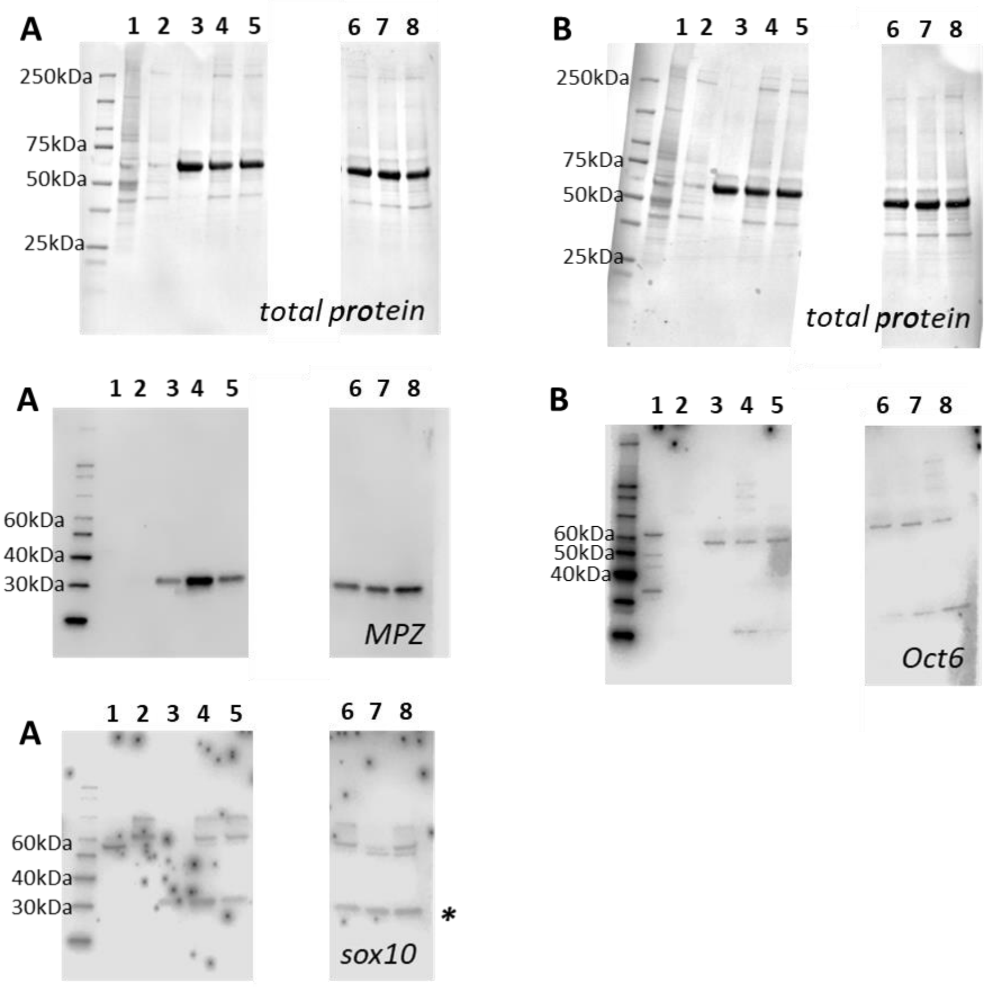


**A**
**1:** brain lysate
**2:** glass
**3:** PLGA 85:15 (co-culture) week 1
**4:** PLGA 50:50 (co-culture) week 1
**5:** PCL (co-culture) week 1
**6:** PCL (co-culture) week 1
**7:** PLGA 85:15 (co-culture) week 1
**8:** PLGA 50:50 (co-culture) week 1

**B**
**1:** brain lysate
**2:** glass
**3:** PLGA 85:15 (co-culture) week 1
**4:** PLGA 50:50 (co-culture) week 1
**5:** PCL (co-culture) week 1
**6:** PCL (co-culture) week 1
**7:** PLGA 85:15 (co-culture) week 1
**8:** PLGA 50:50 (co-culture) week 1

**Figure S1:** Western blots of attachment proteins from co-cultures on polymer films at week 1 (biological replicate 1). Top row: total protein; second row: (A) MPZ, (B) Oct6; third row: (A) sox10; fourth row: sample labels. * Indicates a signal that was not completely removed during blot stripping.

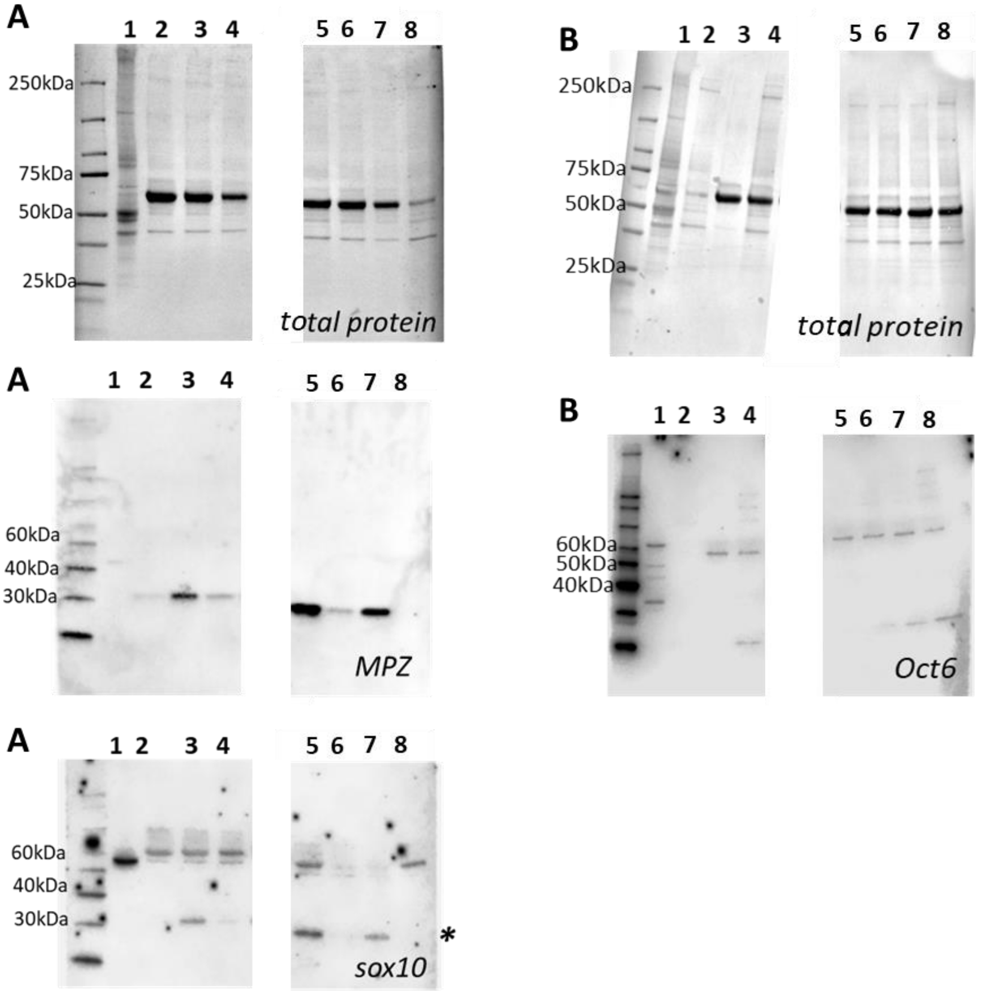


**A**

**1:** brain lysate
**2:** PLGA 85:15 (co-culture) week 1
**3:** PLGA 50:50 (co-culture) week 1
**4:** PCL (co-culture) week 1
**5:** PCL (co-culture) week 1
**6:** PLGA 85:15 (co-culture) week 1
**7:** PLGA 50:50 (co-culture) week 1
**8:** glass

**B**

**1:** brain lysate
**2:** PLGA 85:15 (co-culture) week 1
**3:** PLGA 50:50 (co-culture) week 1
**4:** PCL (co-culture) week 1
**5:** PCL (co-culture) week 1
**6:** PLGA 85:15 (co-culture) week 1
**7:** PLGA 50:50 (co-culture) week 1
**8:** glass

**Figure S2**: Western blots of attachment proteins from co-cultures on polymer films at week 1 (biological replicate 2). Top row: total protein; second row: (A) MPZ, (B) Oct6; third row: (A) sox10; fourth row: sample labels. * Indicates a signal that was not completely removed during blot stripping.

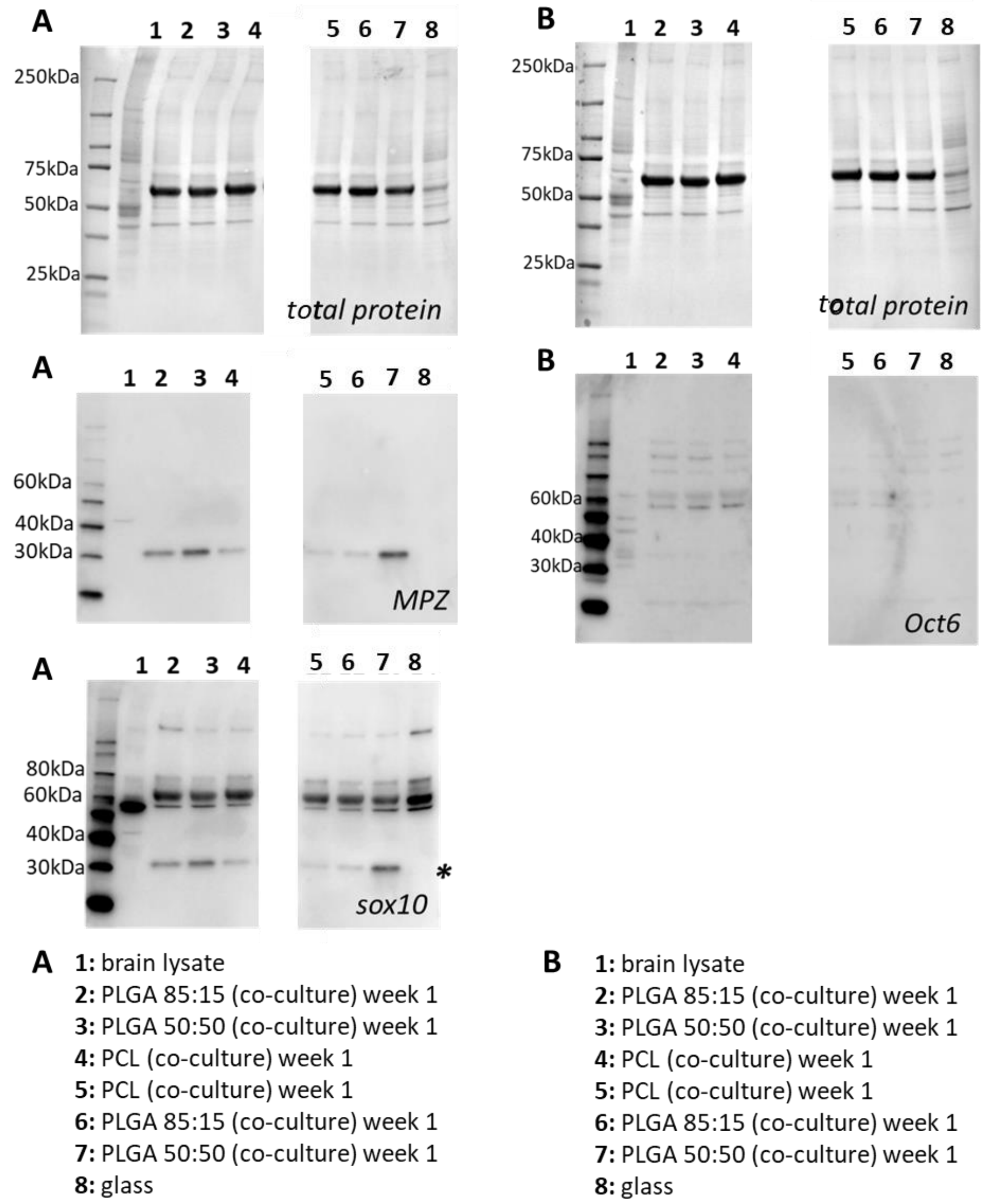


**Figure S3:** Western blots of attachment proteins from co-cultures on polymer films at week 1 (biological replicate 3). Top row: total protein; second row: (A) MPZ, (B) Oct6; third row: (A) sox10; fourth row: sample labels. * Indicates a signal that was not completely removed during blot stripping.

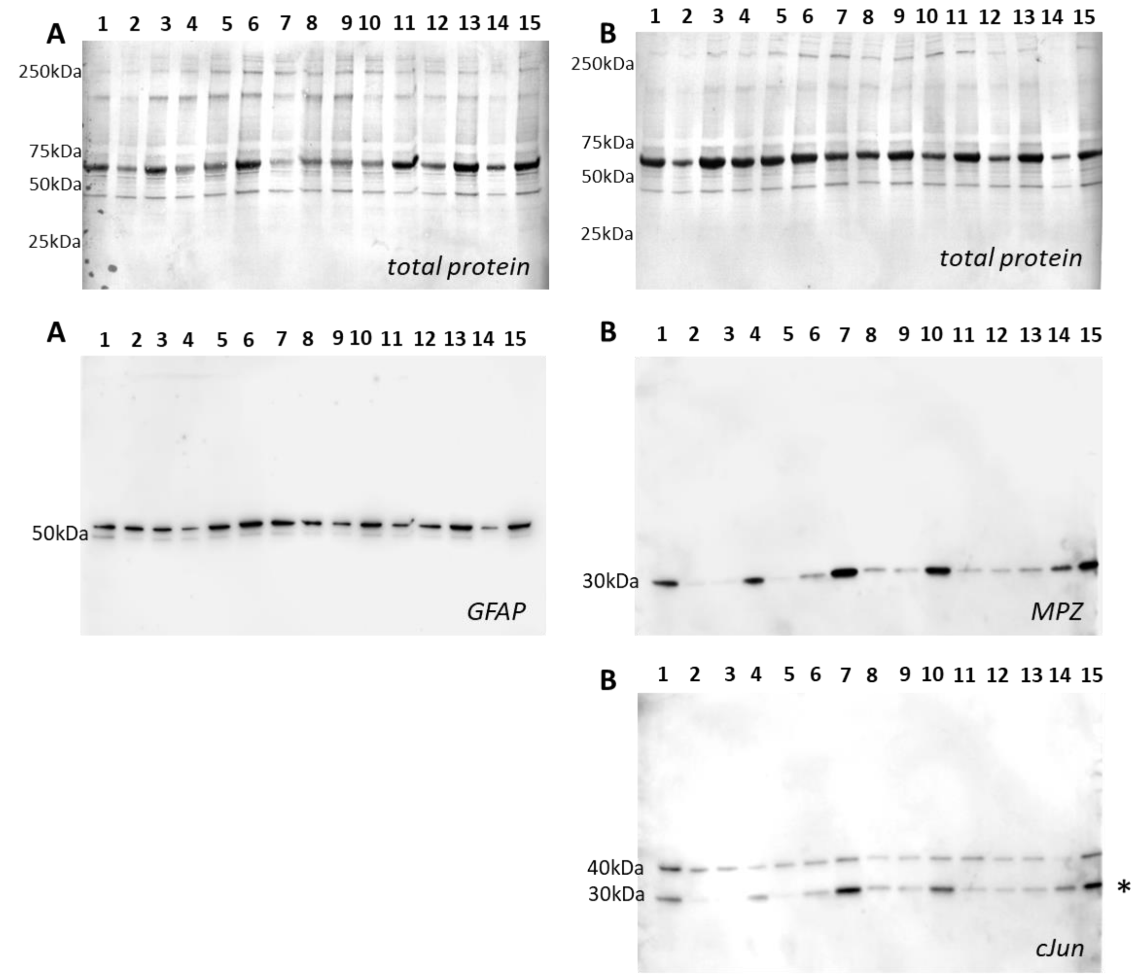


**A**

**1:** PCL (co-culture) week 1
**2:** PCL(co-culture) week 1
**3:** PCL (co-culture) week 1
**4:** PLGA 85:15 (co-culture) week 1
**5:** PLGA 85:15 (co-culture) week 1
**6:** PLGA 85:15 (co-culture) week 1
**7:** PLGA 50:50 (co-culture) week 1
**8:** PLGA 50:50 (co-culture) week 1
**9:** PLGA 50:50 (co-culture) week 1
**10:** PCL (co-culture) week 1
**11:** PCL (co-culture) week 1
**12:** PLGA 85:15 (co-culture) week 1
**13:** PLGA 85:15 (co-culture) week 1
**14:** PLGA 50:50 (co-culture) week 1
**15:** PLGA 50:50 (co-culture) week 1

**B**

**1:** PCL (co-culture) week 1
**2:** PCL(co-culture) week 1
**3:** PCL (co-culture) week 1
**4:** PLGA 85:15 (co-culture) week 1
**5:** PLGA 85:15 (co-culture) week 1
**6:** PLGA 85:15 (co-culture) week 1
**7:** PLGA 50:50 (co-culture) week 1
**8:** PLGA 50:50 (co-culture) week 1
**9:** PLGA 50:50 (co-culture) week 1
**10:** PCL (co-culture) week 1
**11:** PCL (co-culture) week 1
**12:** PLGA 85:15 (co-culture) week 1
**13:** PLGA 85:15 (co-culture) week 1
**14:** PLGA 50:50 (co-culture) week 1
**15:** PLGA 50:50 (co-culture) week 1

**Figure S4**: Western blots of attachment proteins from co-cultures on polymer films at week 1 (all biological replicates). Top row: total protein; second row: (A) GFAP, (B) MPZ; third row: (B) cJun; fourth row: sample labels. * Indicates a signal that was not completely removed during blot stripping.

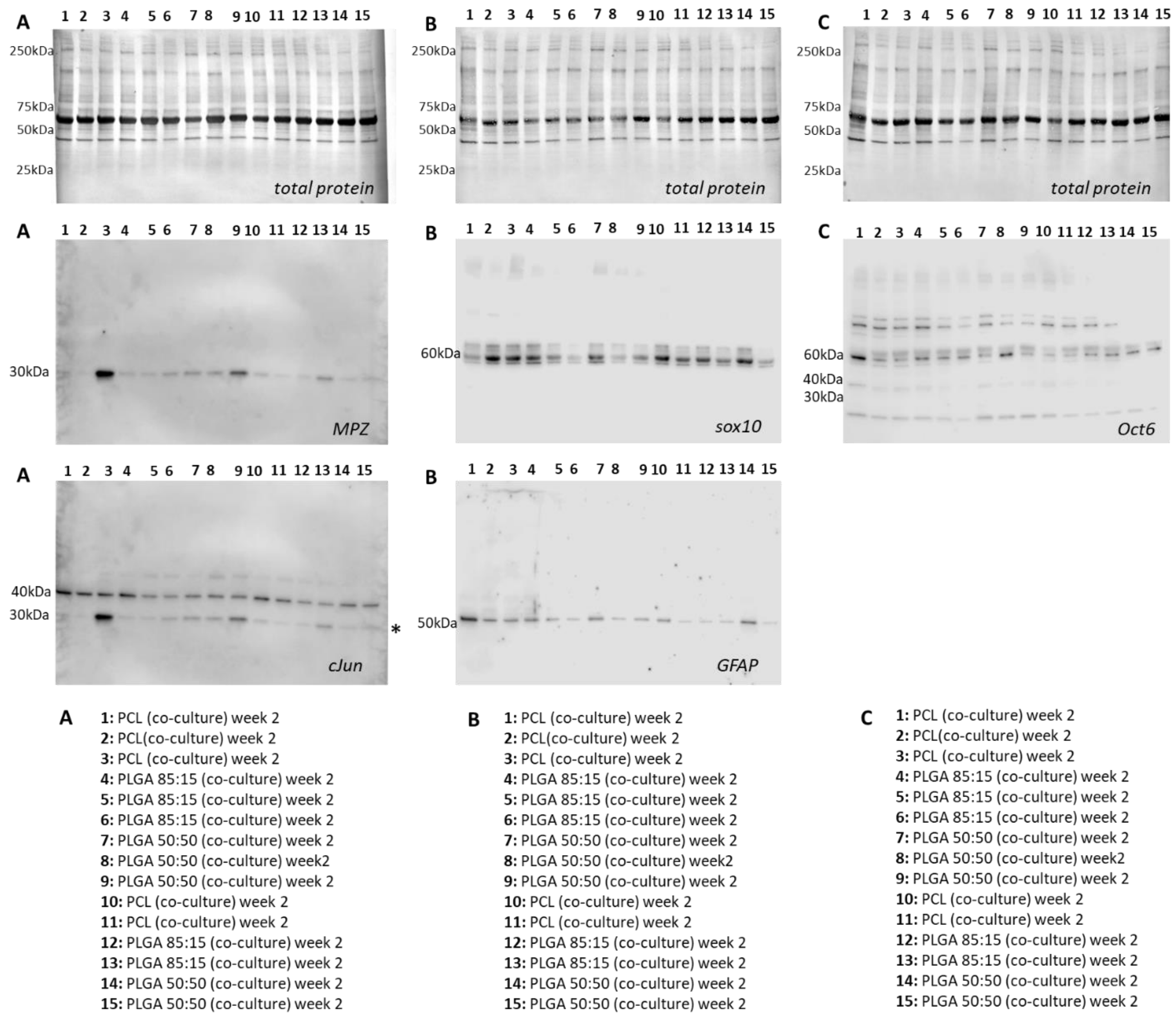


**Figure S5:** Western blots of attachment proteins from co-cultures on polymer films at week 2 (all biological replicates). Top row: total protein; second row: (A) MPZ, (B) sox10, (C) Oct6 staining; third row: (A) cJun, (B) GFAP; fourth row: sample labels. * Indicates a signal that was not completely removed during blot stripping.

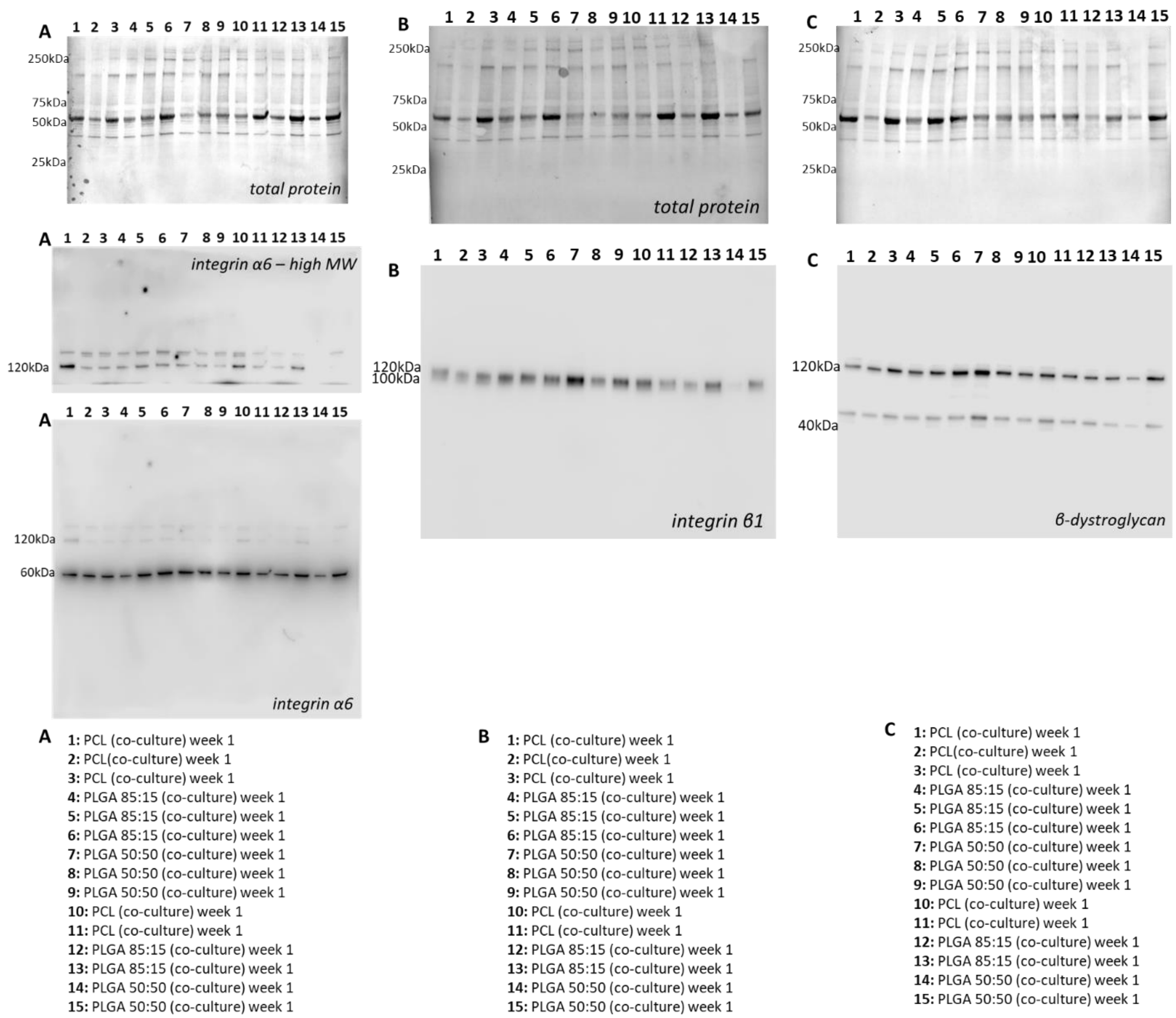


**Figure S6:** Western blots of attachment proteins from co-cultures on polymer films at week 1 (all biological replicates). Top row: total protein; second row: (A) integrin $\alpha_6$ high molecular weight, (B) integrin $\beta_1$, (C) β-dystroglycan staining; third row: (A) integrin $\alpha_6$ full blot, fourth row: sample labels.

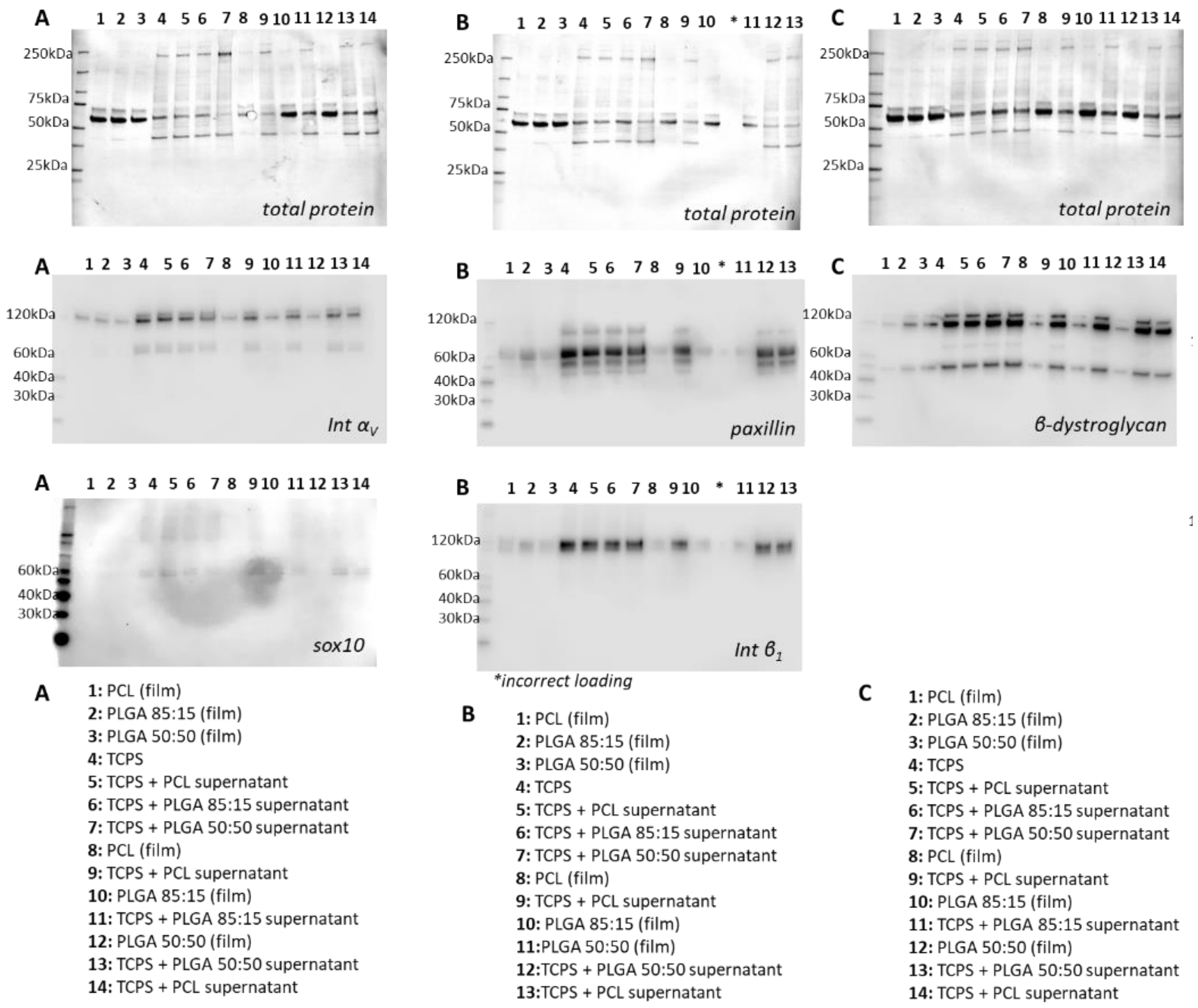


**Figure S7:** Western blots of glial attachment on polymer films (biological replicate 1). Top row: total protein; second row: (A) integrin $\alpha_V$, (B) paxillin, (C) β-dystroglycan staining; third row: (A) sox10, (B) integrin $\beta_1$ staining; fourth row: sample labels. * Indicates a lane with incorrect protein loading.

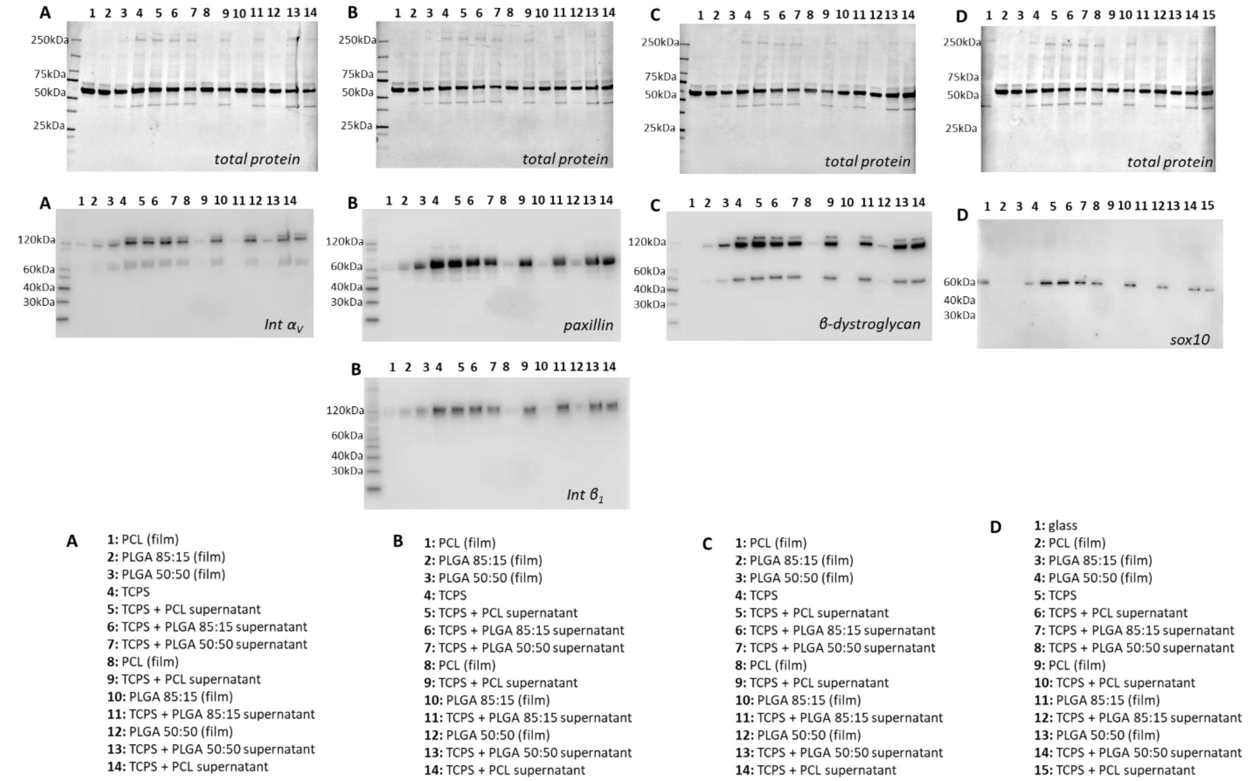


**Figure S8:** Western blots of glial attachment on polymer films (biological replicate 2). Top row: total protein; second row: (A) integrin $\alpha_V$, (B) paxillin, (C) β-dystroglycan, (D) sox10 staining; third row: (B) integrin $\beta_1$ staining; fourth row: sample labels.

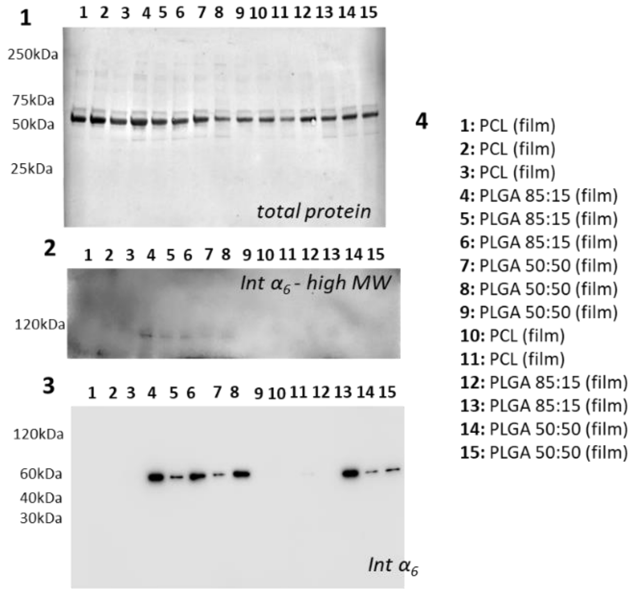


**Figure S9:** Western blot of integrin $\alpha_6$ for glial attachment on polymer films (biological replicates 1-3). (1) total protein; (2) integrin $\alpha_6$ high molecular weight bands only; (3) integrin $\alpha_6$ staining; (4) sample labels.

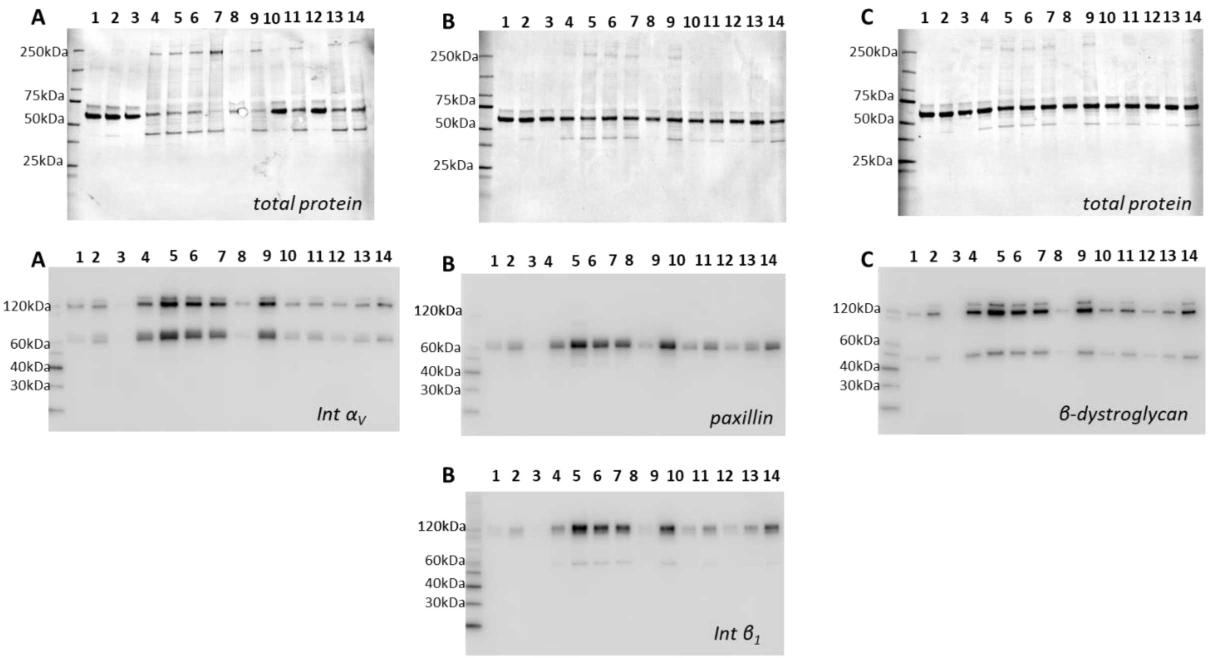


**A**

**1:** PCL (film)
**2:** PLGA 85:15 (film)
**3:** PLGA 50:50 (film)
**4:** TCPS
**5:** TCPS + PCL supernatant
**6:** TCPS + PLGA 85:15 supernatant
**7:** TCPS + PLGA 50:50 supernatant
**8:** PCL (film)
**9:** TCPS + PCL supernatant
**10:** PLGA 85:15 (film)
**11:** TCPS + PLGA 85:15 supernatant
**12:** PLGA 50:50 (film)
**13:** TCPS + PLGA 50:50 supernatant
**14:** TCPS + PCL supernatant

**B**

**1:** PCL (film)
**2:** PLGA 85:15 (film)
**3:** PLGA 50:50 (film)
**4:** TCPS
**5:** TCPS + PCL supernatant
**6:** TCPS + PLGA 85:15 supernatant
**7:** TCPS + PLGA 50:50 supernatant
**8:** PCL (film)
**9:** TCPS + PCL supernatant
**10:** PLGA 85:15 (film)
**11:** TCPS + PLGA 85:15 supernatant
**12:** PLGA 50:50 (film)
**13:** TCPS + PLGA 50:50 supernatant
**14:** TCPS + PCL supernatant

**C**

**1:** PCL (film)
**2:** PLGA 85:15 (film)
**3:** PLGA 50:50 (film)
**4:** TCPS
**5:** TCPS + PCL supernatant
**6:** TCPS + PLGA 85:15 supernatant
**7:** TCPS + PLGA 50:50 supernatant
**8:** PCL (film)
**9:** TCPS + PCL supernatant
**10:** PLGA 85:15 (film)
**11:** TCPS + PLGA 85:15 supernatant
**12:** PLGA 50:50 (film)
**13:** TCPS + PLGA 50:50 supernatant
**14:** TCPS + PCL supernatant

**Figure S10:** Western blots of glial attachment on polymer films (biological replicate 3). Top row: total protein; second row: (A) integrin $\alpha_V$, (B) paxillin, (C) β-dystroglycan staining; third row: (A) sox10, (B) integrin $\beta_1$ staining; fourth row: sample labels. * Indicates a lane with incorrect protein loading.

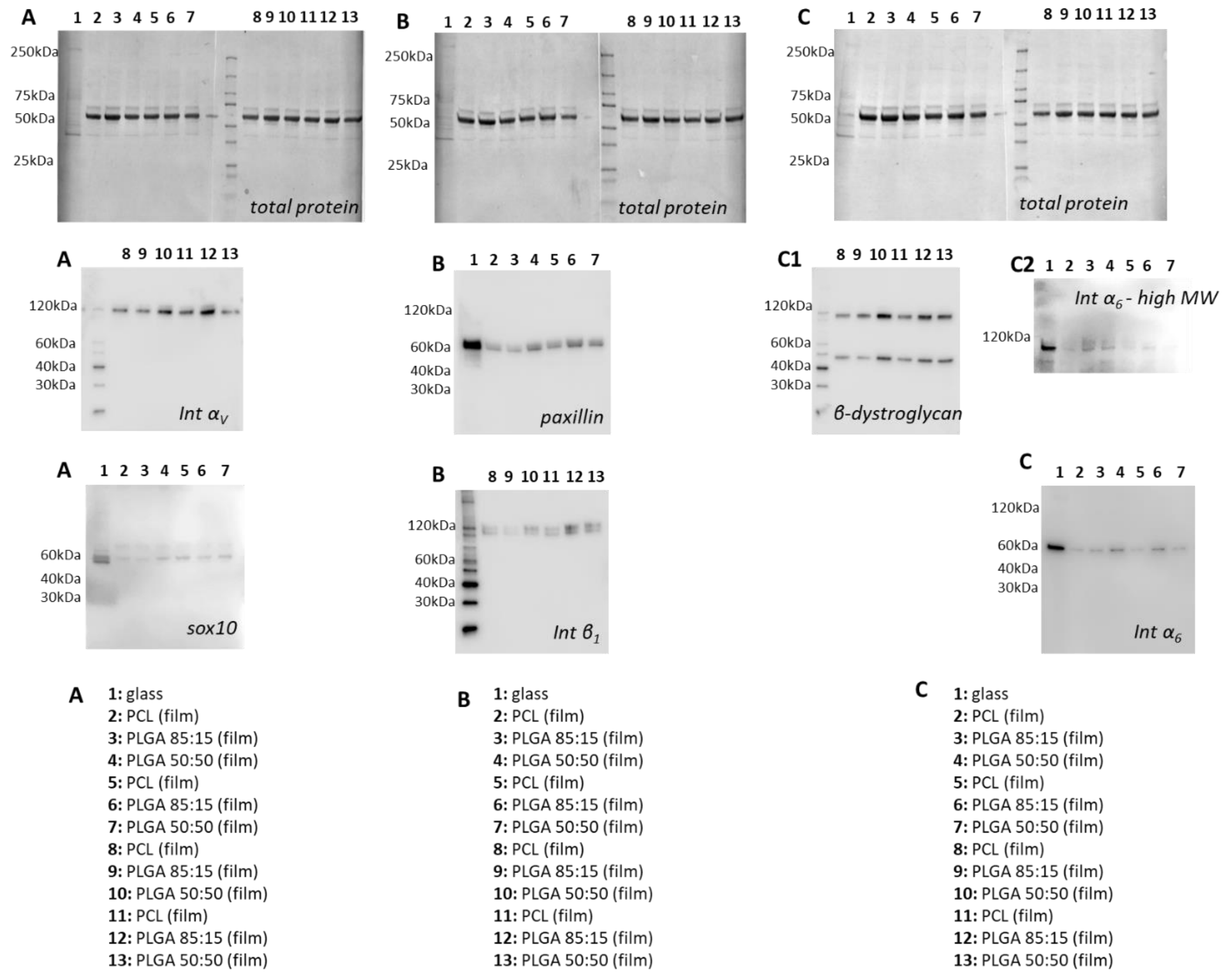


**Figure S11:** Western blots of glial attachment on polymer films (biological replicate 4). Top row: total protein; second row: (A) integrin $\alpha_V$, (B) paxillin, (C1) β-dystroglycan (C2) integrin $\alpha_6$ staining – high molecular weight bands only; third row: (A) sox10, (B) integrin $\beta_1$, (C) integrin $\alpha_6$ staining; fourth row: sample labels.

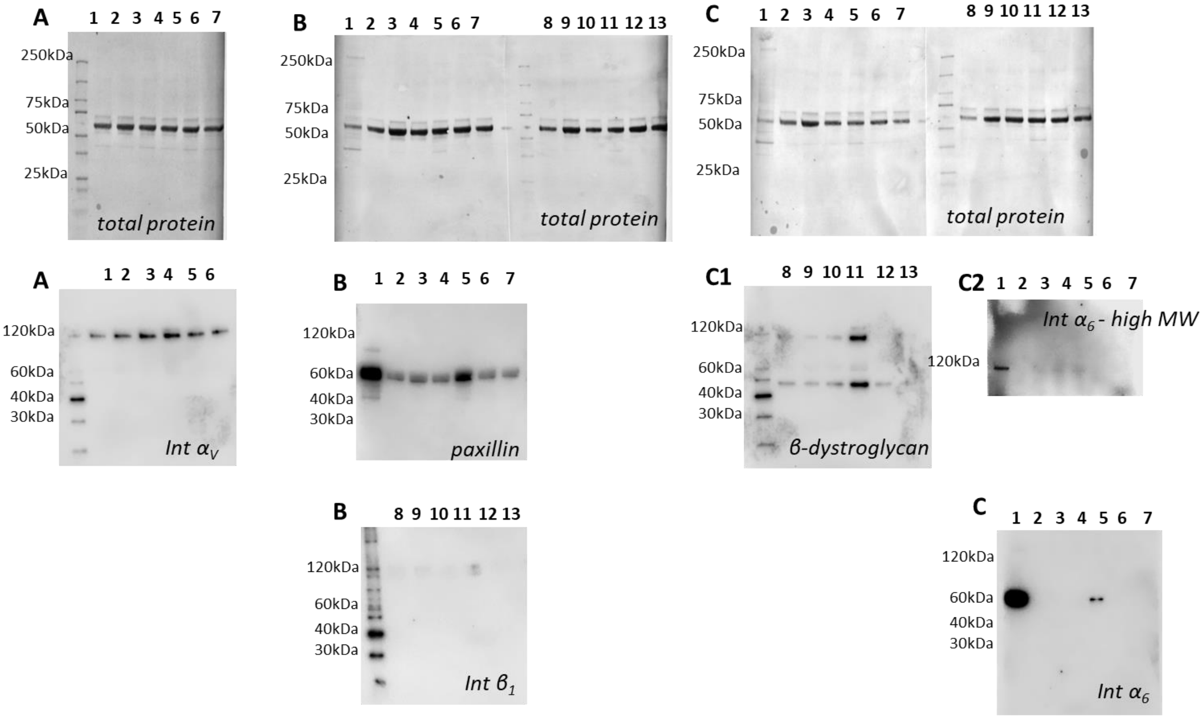


**A**

**1:** PCL (film)
**2:** PLGA 85:15 (film)
**3:** PLGA 50:50 (film)
**4:** PCL (film)
**5:** PLGA 85:15 (film)
**6:** PLGA 50:50 (film)

**B**

**1:** glass
**2:** PCL (film)
**3:** PLGA 85:15 (film)
**4:** PLGA 50:50 (film)
**5:** PCL (film)
**6:** PLGA 85:15 (film)
**7:** PLGA 50:50 (film)
**8:** PCL (film)
**9:** PLGA 85:15 (film)
**10:** PLGA 50:50 (film)
**11:** PCL (film)
**12:** PLGA 85:15 (film)
**13:** PLGA 50:50 (film)

**C**

**1:** glass
**2:** PCL (film)
**3:** PLGA 85:15 (film)
**4:** PLGA 50:50 (film)
**5:** PCL (film)
**6:** PLGA 85:15 (film)
**7:** PLGA 50:50 (film)
**8:** PCL (film)
**9:** PLGA 85:15 (film)
**10:** PLGA 50:50 (film)
**11:** PCL (film)
**12:** PLGA 85:15 (film)
**13:** PLGA 50:50 (film)

**Figure S12:** Western blots of glial attachment on polymer films (biological replicate 5). Top row: total protein; second row: (A) integrin $\alpha_V$, (B) paxillin, (C1) β-dystroglycan (C2) integrin $\alpha_6$ staining – high molecular weight bands only; third row: (B) integrin $\beta_1$, (C) integrin $\alpha_6$ staining; fourth row: sample labels.

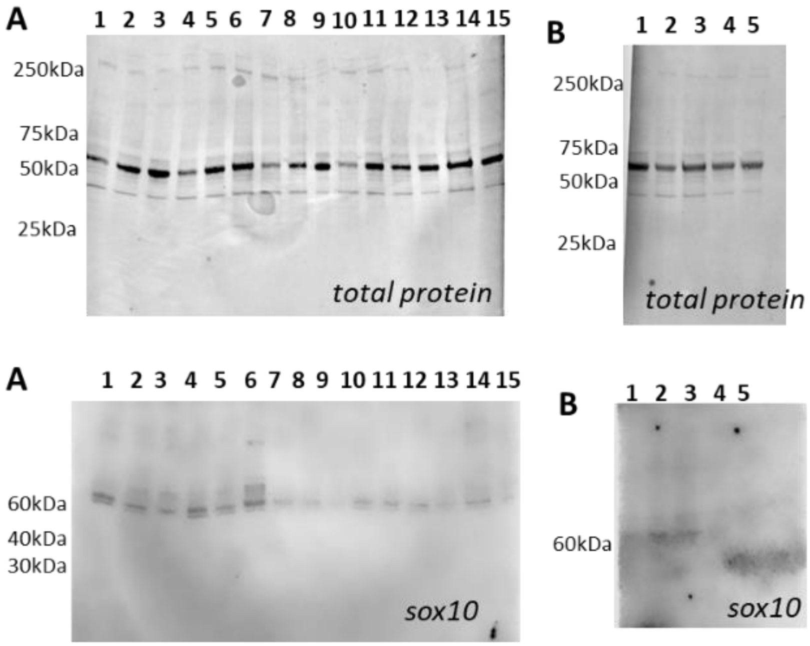


**A**

**1:** TCPS
**2:** TCPS
**3:** TCPS
**4:** TCPS + PCL supernatant
**5:** TCPS + PCL supernatant
**6:** TCPS + PCL supernatant
**7:** TCPS + PLGA 50:50 supernatant
**8:** TCPS + PLGA 50:50 supernatant
**9:** TCPS + PLGA 50:50 supernatant
**10:** TCPS
**11:** TCPS
**12:** TCPS + PCL supernatant
**13:** TCPS + PCL supernatant
**14:** TCPS + PLGA 50:50 supernatant
**15:** TCPS + PLGA 50:50 supernatant

**B**

**1:** TCPS
**2:** TCPS
**3:** TCPS + PLGA 85:15 supernatant
**4:** TCPS + PLGA 85:15 supernatant
**5:** TCPS + PLGA 85:15 supernatant

**Figure S13:** Western blots of glial expression after exposure to degradation products (biological replicates 1-3). Top row: total protein; second row: sox10 staining; third row: sample labels.

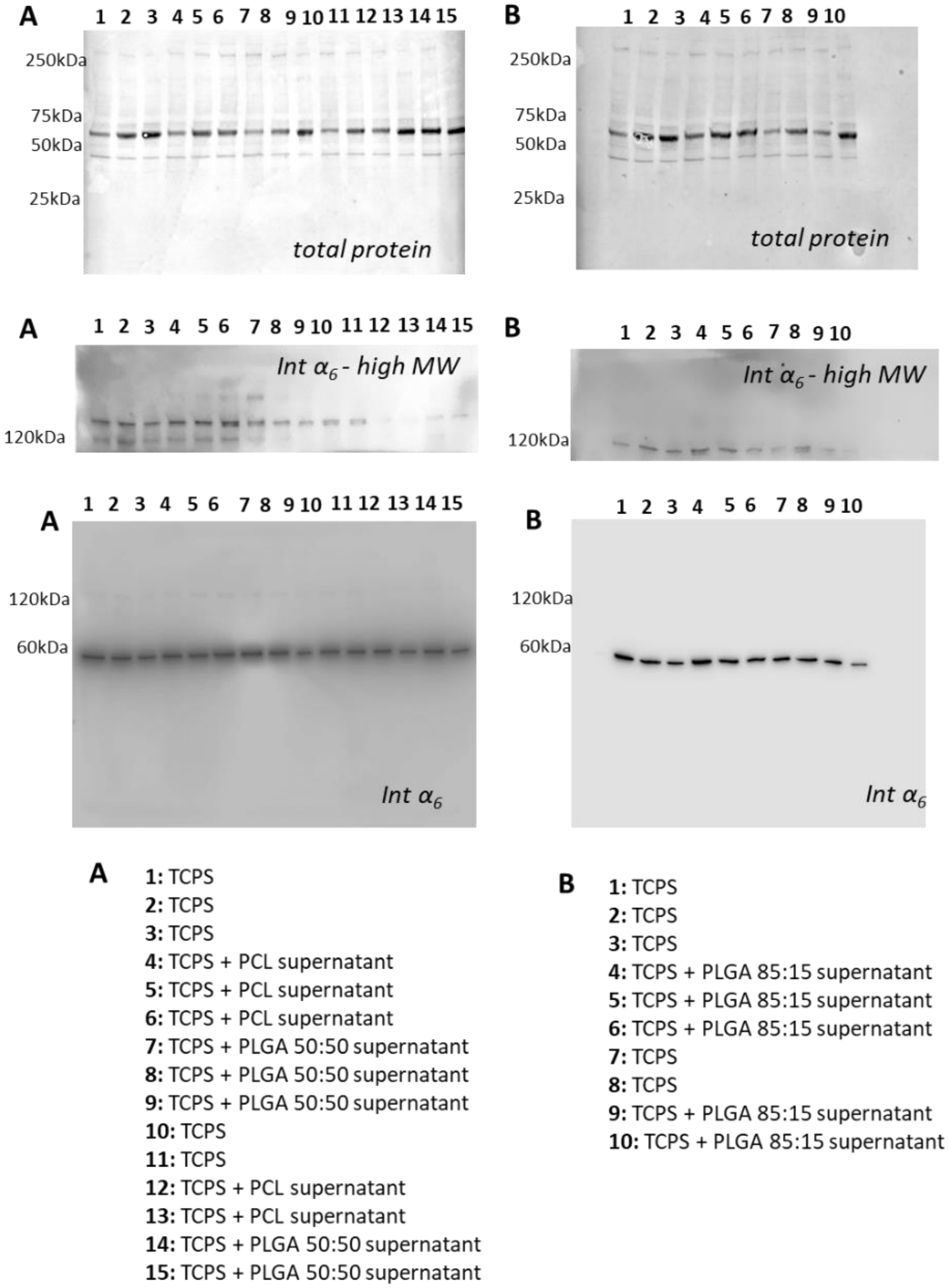


**Figure S14:** Western blots of glial expression after exposure to degradation products (biological replicates 1-3). Top row: total protein; second row: integrin $\alpha_6$ staining – high molecular weight bands only; third row: integrin $\alpha_6$ staining; fourth row: sample labels.

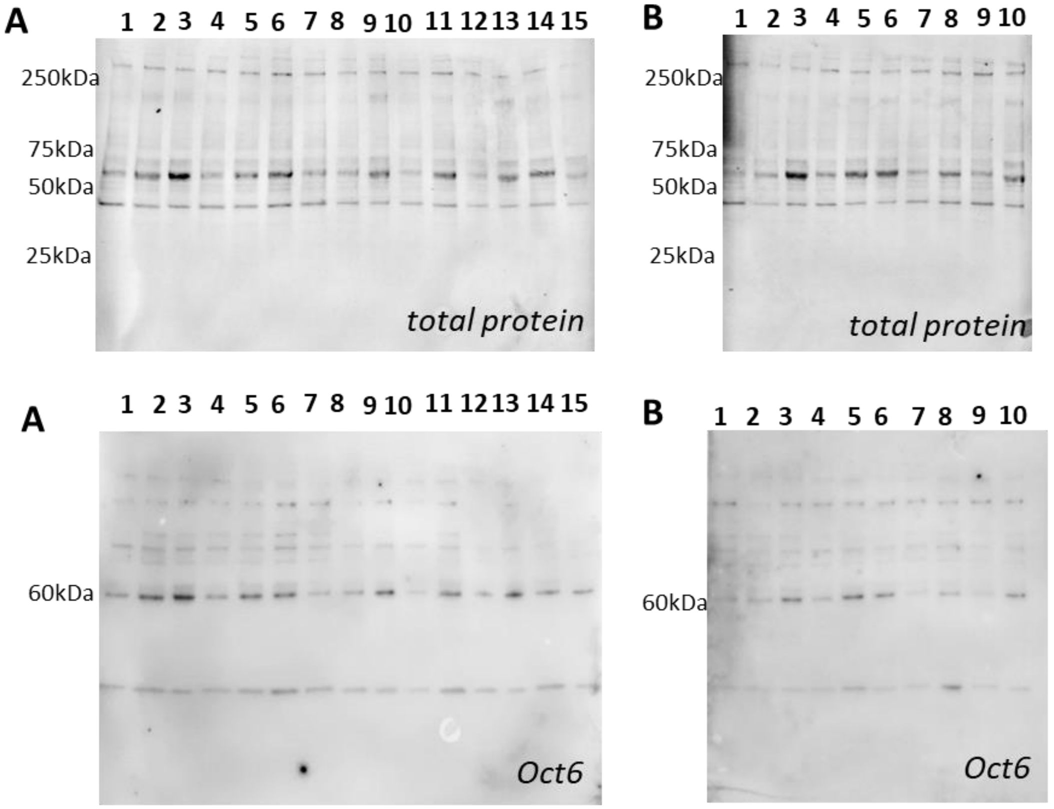


**A**

**1:** TCPS
**2:** TCPS
**3:** TCPS
**4:** TCPS + PCL supernatant
**5:** TCPS + PCL supernatant
**6:** TCPS + PCL supernatant
**7:** TCPS + PLGA 50:50 supernatant
**8:** TCPS + PLGA 50:50 supernatant
**9:** TCPS + PLGA 50:50 supernatant
**10:** TCPS
**11:** TCPS
**12:** TCPS + PCL supernatant
**13:** TCPS + PCL supernatant
**14:** TCPS + PLGA 50:50 supernatant
**15:** TCPS + PLGA 50:50 supernatant

**B**

**1:** TCPS
**2:** TCPS
**3:** TCPS
**4:** TCPS + PLGA 85:15 supernatant
**5:** TCPS + PLGA 85:15 supernatant
**6:** TCPS + PLGA 85:15 supernatant
**7:** TCPS
**8:** TCPS
**9:** TCPS + PLGA 85:15 supernatant
**10:** TCPS + PLGA 85:15 supernatant

**Figure S15:** Western blots of glial expression after exposure to degradation products (biological replicates 1-3). Top row: total protein; second row: Oct6 staining; third row: sample labels.
